# Impacts of an urban sanitation intervention on fecal indicators and the prevalence of human fecal contamination in Mozambique

**DOI:** 10.1101/2021.02.19.432000

**Authors:** David A. Holcomb, Jackie Knee, Drew Capone, Trent Sumner, Zaida Adriano, Rassul Nalá, Oliver Cumming, Joe Brown, Jill R. Stewart

**Affiliations:** Department of Environmental Sciences and Engineering, Gillings School of Global Public Health, University of North Carolina at Chapel Hill, Chapel Hill, North Carolina, United States; Department of Epidemiology, Gillings School of Global Public Health, University of North Carolina at Chapel Hill, Chapel Hill, North Carolina, United States; School of Civil and Environmental Engineering, Georgia Institute of Technology, Atlanta, Georgia, United States; Department of Disease Control, London School of Hygiene & Tropical Medicine, London, United Kingdom; We Consult, Maputo, Mozambique; Instituto Nacional de Saúde, Ministério da Saúde, Maputo, Mozambique

**Keywords:** Diagnostic accuracy, water, sanitation, and hygiene, shared sanitation, microbial source tracking, fecal indicator, qPCR, Bayesian hierarchical model

## Abstract

Fecal source tracking (FST) may be useful to assess pathways of fecal contamination in domestic environments and to estimate the impacts of water, sanitation, and hygiene (WASH) interventions in low-income settings. We measured two non-specific and two human-associated fecal indicators in water, soil, and surfaces before and after a shared latrine intervention from low-income households in Maputo, Mozambique participating in the Maputo Sanitation (MapSan) trial. Up to a quarter of households were impacted by human fecal contamination, but trends were unaffected by improvements to shared sanitation facilities. The intervention reduced *E. coli* gene concentrations in soil but did not impact culturable *E. coli* or the prevalence of human FST markers in a difference-in-differences analysis. Using a novel Bayesian hierarchical modeling approach to account for human marker diagnostic sensitivity and specificity, we revealed a high amount of uncertainty associated with human FST measurements and intervention effect estimates. The field of microbial source tracking would benefit from adding measures of diagnostic accuracy to better interpret findings, particularly when FST analyses convey insufficient information for robust inference. With improved measures, FST could help identify dominant pathways of human and animal fecal contamination in communities and guide implementation of effective interventions to safeguard health.

**SYNOPSIS:** An urban sanitation intervention had minimal and highly uncertain effects on human fecal contamination after accounting for fecal indicator sensitivity and specificity.

**TOC GRAPHIC/ABSTRACT ART:** 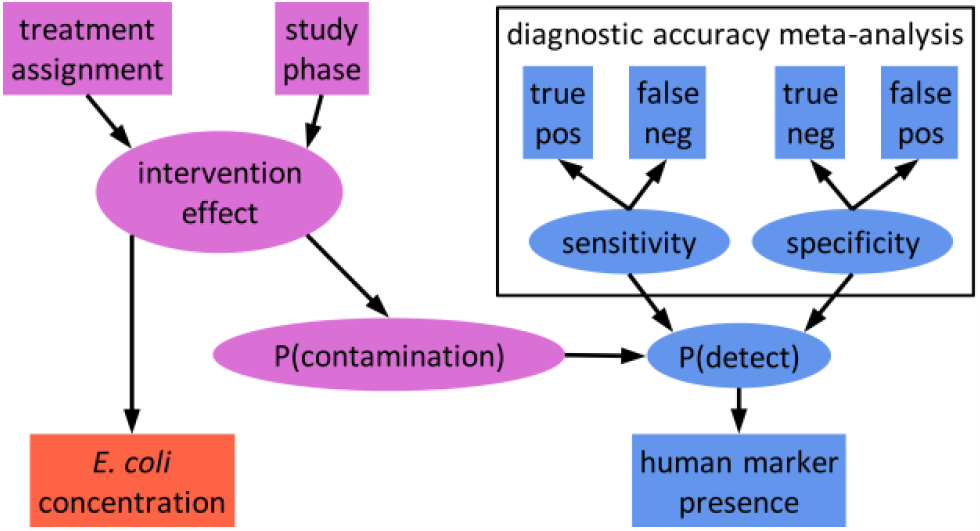

## Introduction

Water, sanitation, and hygiene (WASH) interventions aim to improve health by preventing exposure to enteric pathogens, which are introduced to the environment in the feces of infected human and animal hosts.^1^ Environmental pathways of pathogen exposure include contaminated environmental compartments like water, soil, and surfaces, as well as hands, flies, food, and fomites that have been in contact with contaminated environments.^2–4^ Recent evaluations of a range of WASH interventions found inconsistent and largely negligible impacts on child diarrhea, growth, and enteric infection.^5–12^ Notably, combined interventions did not provide greater protection than their constituent interventions alone, suggesting that key sources of pathogens and pathways of exposure are inadequately addressed by conventional WASH strategies.^6,7,9,13–15^

Characterizing fecal contamination in potential exposure pathways may help explain why specific interventions do or do not improve health by identifying which pathways the intervention interrupts and which remain unaffected. Fecal contamination is typically assessed by measuring fecal indicator organisms, microbes abundant in feces used to infer the presence of fecal contamination and therefore the likely presence of enteric pathogens, which are challenging to measure directly due to their diversity and low environmental concentrations.^15,16^ Indicator organisms can also be used for fecal source tracking (FST) by targeting microbes specific to the feces of a particular host. Animals are important sources of fecal contamination in both domestic and public environments but traditional efforts have focused on preventing exposure to human feces; differentiating between human and various animal feces could inform more appropriate intervention approaches.^4,17–22^

Fecal indicator approaches have increasingly been applied to domestic environments in low-income settings with high burdens of enteric disease.^3,15–18,23–26^ Occurrence of non-specific indicators like *Escherichia coli* is challenging to interpret in these settings due to elevated and highly variable ambient concentrations, possibly from naturalized sources, which are typically assessed in limited numbers of (cross-sectional) observations from each environmental compartment.^16,27–30^ Other than ruminant FST markers, host-associated fecal indicators have demonstrated poor diagnostic accuracy in domestic settings.^17,31,32,26,16^ The use of multiple FST markers has been proposed to help address the limited accuracy of individual indicators.^33,34^ Several studies have calculated the conditional probability of contamination by a specific fecal source given the detection of one or more source-associated indicators in one sample.^31,34–36^ Such analyses provide valuable intuition about the uncertainty associated with individual measurements, which can be particularly important in decision-making contexts like beach closures. To our knowledge, diagnostic performance has not been similarly accounted for when FST has been used to infer patterns and predictors of source-specific fecal contamination in domestic environments, likely overstating the confidence of such estimates.^4,17,18,26,37–39^

In this study, we analyze two non-specific and two human-associated fecal indicators in water, soil, and surfaces from low-income households in Maputo, Mozambique before and after a shared sanitation intervention. We explore the conditional probability of human fecal contamination in individual samples under different prevalence and indicator detection scenarios and develop a Bayesian hierarchical modeling approach that accounts for the diagnostic accuracy of multiple markers to estimate the prevalence of source-specific fecal contamination. Finally, we implement these models using both human markers to estimate intervention effects on the prevalence of human fecal contamination in multiple exposure pathways.

## Materials and Methods

### Study setting and intervention

We characterized fecal contamination of households with children participating in the Maputo Sanitation (MapSan) study (clinicaltrials.gov NCT02362932), a prospective, controlled before and after health impact trial of an urban, onsite sanitation intervention.^40^ The intervention was delivered to compounds (self-defined clusters of households sharing outdoor space) in low-income neighborhoods of Maputo, Mozambique, areas with high burdens of enteric disease and predominantly onsite sanitation infrastructure.^41,42^ Similar compounds that did not receive the intervention were recruited to serve as control sites. At baseline, both intervention and control compounds shared sanitation facilities in poor condition.^26^ The existing shared latrines in intervention compounds were replaced with pour-flush latrines that discharged aqueous effluent to infiltration pits and had sturdy, private superstructures. Intervention latrines were constructed between 2015 – 2016 by the nongovernmental organization (NGO) Water and Sanitation for the Urban Poor (WSUP), which selected intervention sites according to engineering and demand criteria (Table S1).^40^

### Data collection

The intervention impact on fecal contamination was evaluated using a controlled before-and-after (CBA) study design.^5,43^ Intervention compounds were enrolled immediately before the new latrine was opened for use, with concurrent enrollment of control compounds conducted at a similar frequency (Table S1). Follow-up visits to each compound were conducted approximately 12 months following baseline enrollment. We administered compound-, household-, and child-level surveys during both baseline and follow-up visits, as described elsewhere.^5,42^ Concurrent with survey administration during May – August 2015, we opportunistically collected environmental samples at a subset of MapSan study compounds from the shared outdoor space and from each household with children participating in the health study (see Supporting Information [SI]). During the 12-month follow-up phase in June – September 2016, we revisited the original subset of compounds and collected environmental samples from additional study compounds not sampled at baseline, as time permitted.

Detailed descriptions of environmental sample collection, processing, and analysis have been published previously.^26^ Briefly, we assessed fecal indicators in five environmental compartments: compound source water, household stored water, latrine entrance soil, household entrance soil, and household food preparation surfaces (see SI). Source water and latrine soil were sampled once per compound on each visit, while stored water, food preparation surfaces, and household soil were collected from each household with children enrolled in the health impacts study. Samples were processed by membrane filtration, preceded by manual elution for soil and swab samples, and the sample filters were analyzed for microbial indicators of fecal contamination using both culture- and molecular-based detection.^25,26,44^ We enumerated culturable *E. coli* (cEC) from filters on modified mTEC broth (Hi-Media, Mumbai, India) and immediately archived additional filters at -80°C for molecular analysis.^16,45^ Archived filters were analyzed by three locally validated real-time polymerase chain reaction (qPCR) assays targeting fecal microbe genes. The EC23S857 (EC23S) assay targets *E. coli* and served as an indicator of non-specific fecal contamination, while HF183/BacR287 (HF183) and Mnif both target microbes specific to human feces and served as indicators of human-source fecal contamination.^46–48^ Limits of detection for each assay were previously determined using receiver operating characteristic (ROC) analysis to identify optimal quantification cycle (Cq) cutoff values (see SI).^26,49^

DNA was isolated from soil and surface sample filters using the DNeasy PowerSoil Kit (Qiagen, Hilden, Germany) and from water sample filters with the DNA-EZ ST01 Kit (GeneRite, North Brunswick, NJ, USA), with a positive control (PC) and negative extraction control (NEC) included in each batch of up to 22 sample filters. PCs consisted of a clean filter spiked with 2 × 10^8^ copies of each composite DNA standard (Table S4).^26^ Filters were treated with 3 µg salmon testes DNA (MilliporeSigma, Burlington, MA, USA) immediately before extraction as a specimen processing control (SPC) to assess PCR inhibition.^50,51^ We tested each extract with four qPCR assays using a CFX96 Touch thermocycler (Bio-Rad, Hercules, CA), three targeting fecal microbes and Sketa22 targeting the salmon DNA SPC, with 10% of each sample type analyzed in duplicate for all microbial targets.^52^ Each reaction consisted of 12.5 µL TaqMan Environmental Master Mix 2.0, 2.5 µL 10x primers/probe mix, 5 µL nuclease free water (NFW), and 5 µL DNA template, for 25 µL total reaction volume. After an initial 10-minute, 95°C incubation period, cycling conditions specified by the original developers were followed for each assay (Table S3). Samples with Sketa22 quantification cycle (Cq) values > 3 above the mean Cq of extraction controls (NEC and PC) were considered inhibited and diluted 1:5 for further analysis. Each plate included three no-template controls (NTCs) and five-point, ten-fold dilution series of three extracted PCs, corresponding to triplicate reactions with 10^5^ – 10^1^ or 10^6^ – 10^2^ target gene copies (gc). Target concentrations were estimated from calibration curves fit to the standard dilution series using multilevel Bayesian regression with varying slopes and intercepts by extraction batch and instrument run (see SI).^53^ Fecal indicator concentrations were log_10_ transformed and expressed as log_10_ colony forming units (cfu) or gc per 100 mL water, 100 cm^2^ surface, or 1 dry gram soil.

### Estimating intervention effects

We used a difference-in-differences (DID) approach to estimate the effect of the intervention on fecal indicator occurrence. DID enables unbiased estimation of the treatment effect in the absence of randomization, including when different samples of each group are observed pre- and post-treatment, under the “parallel trend” assumption that all unmeasured time-varying covariates related to the outcome are constant across treatment groups and that unmeasured covariates varying between treatment groups are constant through time.^43,54,55^ Although we estimated gene copy concentrations for all fecal indicators assessed by qPCR, we treated the human markers as binary, diagnostic tests of the presence or absence of human fecal contamination due to their relatively low baseline detection frequency (and limited availability of concentration data as a result).^26^ By contrast, *E. coli* was detected in the large majority of baseline samples by both culture and qPCR approaches; treating such outcomes as presence/absence would discard a great deal of information conveyed by the *E. coli* concentration measurements, producing a binary outcome with very little variation. Direct DID estimates for the mean concentration of non-specific indicators and the prevalence of human-associated indicators were obtained using a bootstrap approach with 2000 samples. We calculated the mean concentration or prevalence in each of the four design strata (pre-treatment intervention compounds, post-treatment intervention compounds, pre-treatment control compounds, and post-treatment controls) by sample type, from which the DID was calculated directly (see SI). Bootstrap 95% compatibility intervals (CI) were obtained as the 2.5 and 97.5 percentile values of the bootstrap samples.^56^

We also conducted regression analyses incorporating potential confounding variables to obtain conditional DID estimates. We used the product-term representation of the DID estimator, in which binary indicators of treatment group, study phase, and their product (interaction) were included as linear predictors. The coefficient on the product term provides the conditional DID estimate.^54,57^ Separate models were fit for each combination of fecal indicator and sample type using Bayesian multilevel models with compound-varying intercepts. Censored linear regression was used to estimate the intervention impact on the log_10_ concentration of non-specific indicators and the effect of the intervention on human-associated indicator prevalence was estimated using logistic regression and the prevalence odds ratio (POR) as the measure of effect.^58,59^ Models were fit with the package **brms** in **R** version 4.0.2 using 1500 warmup and 1000 sampling iterations on four chains (see SI for prior distributions).^58,60^ Estimates of the intervention effect were summarized by the mean and central 95% CI of the resulting 4000 posterior draws.

Adjusted models included variables for selected compound, household, meteorological, and sample characteristics. Compound population density, presence of domestic animals, and asset-based household wealth scores were derived from household and compound surveys administered during each study phase.^42,61^ Previous day mean temperature and seven-day antecedent rainfall were drawn from daily summary records for a local weather station. For stored water samples, we considered whether the storage container was covered and if the mouth was wide enough to admit hands. The surface material was considered for food surface swabs, and for soil samples we accounted for sun exposure and visibly wet soil surfaces. Covariate data sources and processing have been described previously.^26,42^

### Conditional probability analysis

Both HF183 and Mnif were previously found to frequently misdiagnose human feces in our study area.^26^ An indicator’s diagnostic accuracy is described by its sensitivity (Se), the probability of detecting the indicator when contamination is present, and specificity (Sp), the probability of not detecting the indicator when contamination is absent. The probability that a positive sample is contaminated depends on the marker sensitivity and specificity and the prevalence of human fecal contamination. This marginal probability of contamination can be approximated as the frequency of indicator detection among all samples to explore indicator reliability in a specific study.^31^ We assessed the probability that human feces were present in an environmental sample in which HF183 or Mnif was detected using Bayes’ Theorem and the local sensitivity and specificity of the two markers (see SI).^34–36^ We calculated the conditional probability of contamination for HF183 and Mnif separately and for each combination of the two indicators by sample type. The marginal probability of contamination was approximated as the detection frequency of HF183 among all samples of a given type.

### Accounting for diagnostic accuracy

Fecal indicator measurements are used as proxies for unobserved fecal contamination to estimate its prevalence and associations of interest, such as the effects of mitigation practices. This approach is vulnerable to measurement error, illustrated by the limited diagnostic accuracy of many host-associated fecal indicators.^16^ Bias due to inaccurate diagnostic tests can be mitigated by incorporating external information on the sensitivity and specificity of the test.^62^ The expected detection frequency, *p*, of a test with sensitivity *Se* and specificity *Sp* is given by

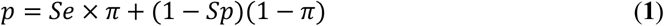

for an underlying condition with prevalence *π*.^62,63^ We adapted the approach of Gelman and Carpenter to estimate the intervention effect on human fecal contamination prevalence from observations of human-associated fecal indicators by incorporating external information on indicator performance within a Bayesian hierarchical framework.^63^ We included the product-term representation of the DID estimator and other covariates as linear predictors of the prevalence log-odds. Assuming indicator detection in the *i*th of *n* samples, *y*_*i*_, was Bernoulli-distributed with probability *p*_*i*_, where *p*_*i*_ was related to the prevalence as shown in Equation (**1**), the accuracy-adjusted prevalence model was

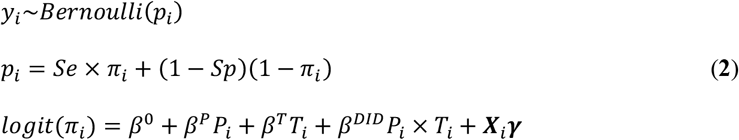

where *β*^0^ is the intercept; *β*^*P*^, *β*^*T*^, and *β*^*DID*^ are the parameters corresponding to indicators for study phase (*P*), treatment group (*T*), and their product; and ***γ*** is a *p* × 1 vector of regression coefficients corresponding to the *p* additional covariates in the *n* × *p* matrix ***X***.

We fit three models that differed by definition of *Se* and *Sp*. In the simplest case (Model 1), we assumed a perfectly accurate test with *Se* = *Sp* = 1, thus *p* = *π*. The second model (Model 2) incorporated observations from the local validation analysis by assuming

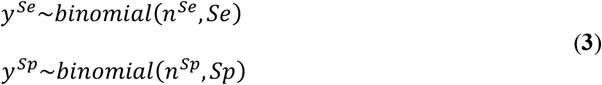

for *y*^*Se*^ positive results in *n*^*Se*^ human fecal samples and *y*^*Sp*^ negative results in *n*^*Sp*^ non-human fecal samples. Because our validation sample set was small and performance estimates vary widely between studies, we fit a third model (Model 3) featuring a meta-analysis of indicator sensitivity and specificity (see SI). We assumed the log-odds of the sensitivity in the *k*th study, *Se*_[*k*]_, were normally distributed with mean *μ*^*Se*^ and SD *σ*^*Se*^, such that

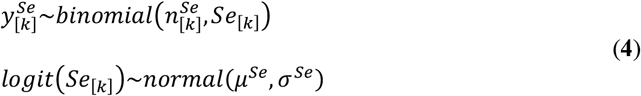

with an equivalent structure for the specificity. We assigned *k* = 1 to our local validation study, using *Se*_[1]_ and *Sp*_[1]_ as the values of *Se* and *Sp* in Equation (**2**).^26,63^ This emphasized the local performance data while allowing information from other settings to influence the estimates through partial pooling, with the extent of pooling learned from the data (expressed through *σ*^*Se*^ and *σ*^*Sp*^).^59^

### Modeling latent human fecal contamination

Fecal contamination can be understood as a latent environmental condition for which fecal indicators serve as imperfect diagnostic tests.^64,65^ Information from multiple fecal indicators may be utilized by modeling each as arising from the same underlying contamination to potentially improve inference. We extended the meta-analytic model (Model 3) to include observations of both HF183 and Mnif in the same samples (Model 4), with separate detection probabilities, 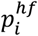 and 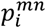, obtained from indicator-specific sensitivity and specificity estimates applied to the same underlying prevalence, *π*_*i*_. As in previous models, the DID estimator and other predictor variables were included in a linear model on the log-odds of *π*_*i*_, assuming that intervention effects and other covariates acted directly on the latent prevalence.

As environmental compartments from the same compound share sources of fecal exposure, we extended the previous model to simultaneously consider observations of latrine soil, household soil, and stored water in each compound (Model 5). Sample type-specific prevalence variables, 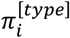, were modeled as linear deviations from a latent compound-level prevalence *π*_*j*_ on the log-odds scale:

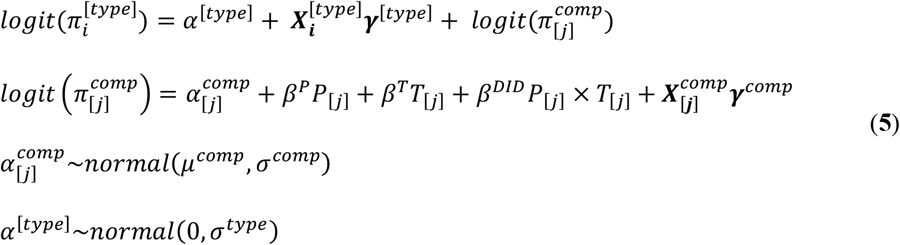

for sample *i* of a given *type* (latrine soil, household soil, or stored water) in compound *j*, where 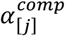 is a compound-varying intercept and *α*^[*type*]^ is a varying intercept by sample type. Compound-level predictors, including the DID estimator terms, were placed on the compound-prevalence log-odds.^63,66^ Parameters for sample-level and meteorological predictors in 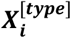 were estimated separately for each sample type.

We coded each model in the probabilistic programming language **Stan** and fit the models using the **RStan** interface with four chains of 1000 warmup and 1000 sampling iterations each, for a total of 4000 posterior samples (see SI for Stan code and discussion of prior distributions).^67,68^ Models 1-3 were fit separately for HF183 and Mnif in each sample type (latrine entrance soil, household entrance soil, and stored water), Model 4 was fit separately to each sample type, and a single Model 5 fit was produced incorporating both indicators and all sample types. In addition to the DID POR given by the product-term parameter, we used the posterior predictive distribution to estimate the prevalence of human fecal contamination in each stratum and to directly calculate DID on the probability scale.^59,69^ Models were adjusted for the same covariates as the DID regression models.

### Ethical approval

This study was approved by the Institutional Review Board of the University of North Carolina at Chapel Hill (IRB # 15-0963) and the associated health study was approved by the Comité Nacional de Bioética para a Saúde (CNBS), Ministério da Saúde, Republic of Mozambique (333/CNBS/14), the Ethics Committee of the London School of Hygiene and Tropical Medicine (reference # 8345), and the Institutional Review Board of the Georgia Institute of Technology (protocol # H15160). Environmental samples were only collected from households with enrolled children for whom written, informed parental or guardian consent had been given.

## Results

### Sample characteristics

We collected a total of 770 environmental samples from 507 unique locations at 139 households in 71 compounds. Samples were collected both pre- and post-intervention at 263 locations (52%), for a total of 526 paired samples and 244 unpaired samples (Table S2). Characteristics expected to confound the relationship between sanitation and fecal contamination were largely similar between treatment arms during each study phase (Table 1). Cumulative precipitation was higher on average in intervention compounds at baseline and in control compounds at follow-up. Water storage containers were also more frequently covered in intervention (75%) than control households (57%) at baseline, though the majority of containers were covered in all strata. Soil surfaces were more often visibly wet in control households (51%) than intervention (33%) at follow-up, both of which were lower than at baseline (57% and 48%, respectively). Most food preparation surfaces were plastic, though more often so in control households during both study phases. A higher percentage of compounds from both treatment arms reported owning domestic animals at follow-up (80–88%) than baseline (47–68%), which may be related to differences in the questionnaire between survey phases. Median household wealth was 40–45 on a 100-point index, with higher variance among controls at follow-up. Median compound population density ranged from 5.5–8.1 residents/100 m^2^.

**Table 1.** Characteristics of Maputo Sanitation study compounds and households selected for environmental sampling, samples collected, and sampling dates, stratified by study phase and treatment arm.

| characteristic | level | metric | before |  |  |  | after |  |  |  |
| --- | --- | --- | --- | --- | --- | --- | --- | --- | --- | --- |
|  |  |  | control |  | intervention |  | control |  | intervention |  |
|  |  |  | N | summary | N | summary | N | summary | N | summary |
| animals present | compound | n (%) | 32 | 15 (47) | 25 | 17 (68) | 30 | 24 (80) | 34 | 30 (88) |
| population density (persons/100 m<sup>2</sup>) | compound | median (IQR<sup>a</sup>) | 29 | 5.5 (3.5) | 23 | 8.1 (5.9) | 28 | 5.9 (4.8) | 33 | 6.7 (4.6) |
| wealth index (0 - 100) | household | median (IQR) | 51 | 43 (12) | 40 | 43 (12) | 55 | 45 (19) | 52 | 44 (14) |
| previous day mean temperature (°C) | date | median (IQR) | 19 | 21 (2) | 16 | 20 (2) | 17 | 20 (1) | 17 | 21 (3) |
| seven-day cumulative precipitation (mm) | date | median (IQR) | 19 | 9 (3) | 16 | 14 (3) | 17 | 13 (39) | 17 | 7 (0) |
| water container covered | sample | n (%) | 44 | 25 (57) | 28 | 21 (75) | 38 | 21 (55) | 47 | 30 (64) |
| narrow-mouth water container | sample | n (%) | 44 | 13 (30) | 28 | 10 (36) | 38 | 13 (34) | 47 | 14 (30) |
| plastic food surface material | sample | n (%) | 34 | 30 (88) | 23 | 18 (78) | 29 | 26 (90) | 36 | 29 (81) |
| shaded latrine soil | sample | n (%) | 32 | 24 (75) | 17 | 12 (71) | 30 | 25 (83) | 30 | 22 (73) |
| shaded household soil | sample | n (%) | 42 | 31 (74) | 28 | 24 (86) | 35 | 32 (91) | 39 | 31 (79) |
| wet latrine soil surface | sample | n (%) | 32 | 20 (62) | 17 | 13 (76) | 30 | 18 (60) | 30 | 21 (70) |
| wet household soil surface | sample | n (%) | 42 | 24 (57) | 27 | 13 (48) | 35 | 18 (51) | 39 | 13 (33) |
| latrine soil moisture (%) | sample | median (IQR) | 33 | 9.8 (9.8) | 23 | 8.4 (7.2) | 30 | 10.0 (7.9) | 30 | 8.7 (8.3) |
| household soil moisture (%) | sample | median (IQR) | 49 | 9.9 (8.6) | 35 | 6.9 (6.1) | 47 | 7.8 (5.4) | 43 | 5.4 (5.9) |

<sup>a</sup> interquartile range

### Fecal indicator occurrence

At least one fecal indicator was detected in 94% of samples (720/770) and *E. coli* was detected in 718 samples: by culture in 81% (611/755) and by qPCR in 86% (655/763). Mean cEC concentrations were lower at follow-up for all sample types in both treatment arms, a pattern not observed for EC23S concentrations (Figure 1). Of the 763 samples tested for human-associated indicators, 28% (217) were positive for at least one human marker. Human-associated indicators were common in soils (23–65% prevalence, across treatment groups and study phases) but only HF183 was regularly detected in stored water (10–22%) and both indicators were rare on food surfaces (0–9%). qPCR calibration curves (Table S5), detection limits (Table S6), and the results of laboratory quality controls are presented in the SI.

**Figure 1.**
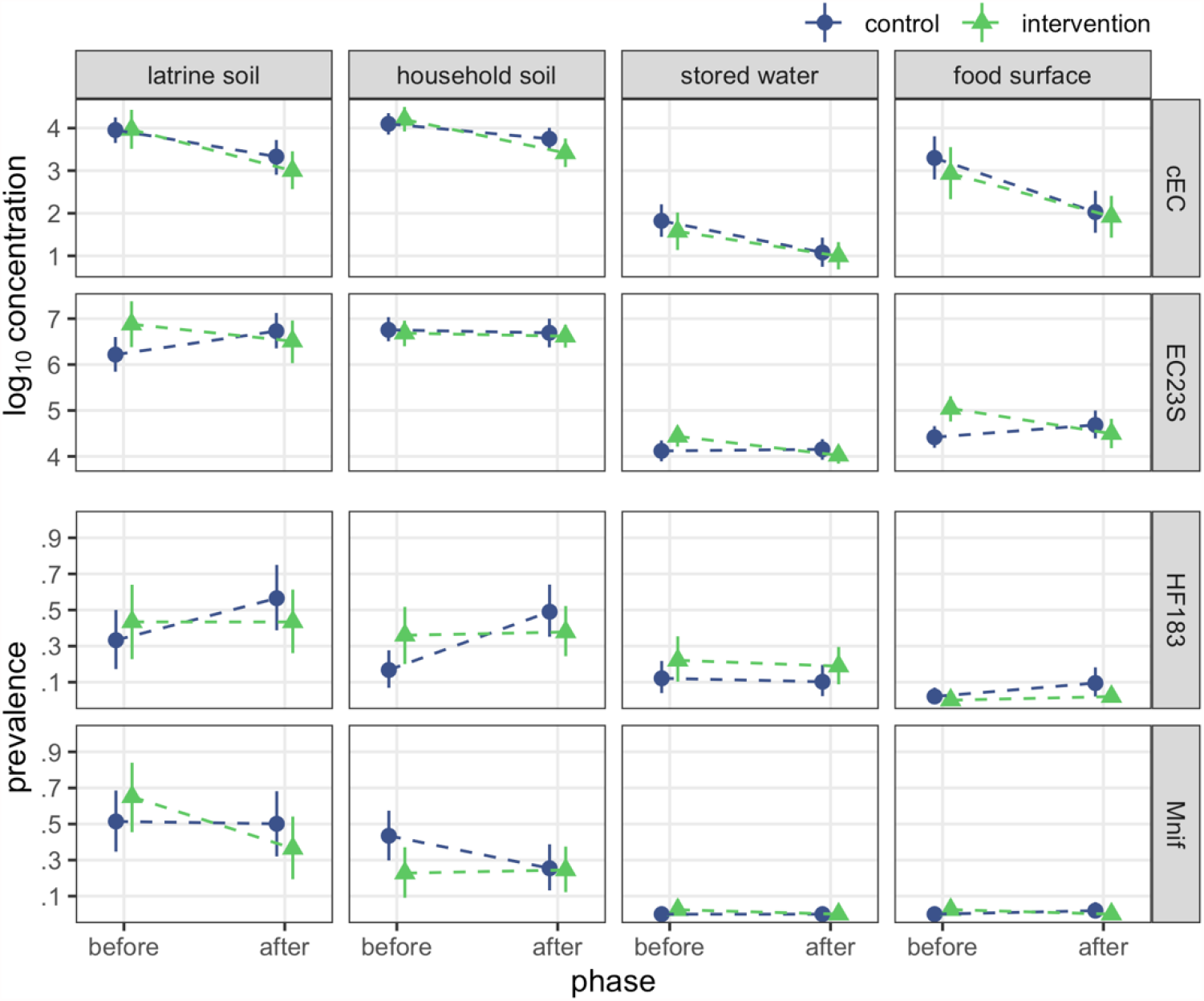
Bootstrap estimates of fecal indicator occurrence by study phase and treatment arm. Points indicate mean log_10_ concentration for *E. coli* indicators and prevalence of human-associated indicators, with bars presenting bootstrap 95% CIs.

Bootstrap DID estimates suggest the intervention reduced EC23S concentrations on food preparation surfaces and HF183 prevalence in household soil but minimally impacted fecal indicator occurrence in other sample types (Table S7). Notably, HF183 prevalence in household soil was similar among intervention households in both study phases but increased among control compounds at follow-up. By contrast, model-based DID estimates, adjusted for potential confounding, were consistent with no intervention effect on food preparation surface EC23S concentration or household soil HF183 prevalence (Table S8). Adjusted models instead indicate the intervention reduced latrine soil concentrations of EC23S [mean difference: -1.2 (95% CI: - 2.1, -0.30) log_10_ gc/dry g]. Although several sample characteristics were imbalanced between treatment arms and study phases (Table 1), estimates from models that adjusted for these variables were largely similar to the unadjusted models, with adjusted estimates marginally closer to the null in most cases (Table S8). EC23S concentrations in latrine soil were again the exception, with a substantially larger reduction obtained under the adjusted model than the unadjusted estimate of -0.84 (95% CI: -1.6, -0.02) log_10_ gc/dry g. Due to low detection frequency, models were not fit for either human marker on food surfaces or for Mnif in stored water; source water samples were excluded from all analyses.^26^

### Conditional probability of human fecal contamination

The probability that a sample is contaminated with human feces given the detection of a human indicator is a function of the indicator’s sensitivity and specificity (Table S9) and the prevalence of human contamination in the study environment. At 15% prevalence (approximately the detection frequency of HF183 in stored water), the probability of human contamination given a positive test was 26% for HF183 and 30% for Mnif. Only with prevalence above 30–35% was detecting either indicator more likely than not to correctly diagnose human fecal contamination. Combining test results from both indicators improved identification of human contamination, increasing the probability of contamination to 45% when both markers were positive and the prevalence was 15% (Figure 2). However, the two human markers frequently disagreed when assessed in the same sample, conflicting in 44% of household soil, 43% of latrine soil, and 15% of stored water samples. Furthermore, at 44% prevalence (the highest detection frequency for HF183, observed in latrine soils), there remained a >20% chance that a sample positive for both indicators was not contaminated. Among lower-prevalence sample types the conditional probability never reached 50%. Unless the background prevalence in the study area was about 45% or greater, it is unlikely that the use of HF183 and Mnif reliably identified human contamination in individual samples, particularly given the frequent disagreement between the two markers.

**Figure 2.**
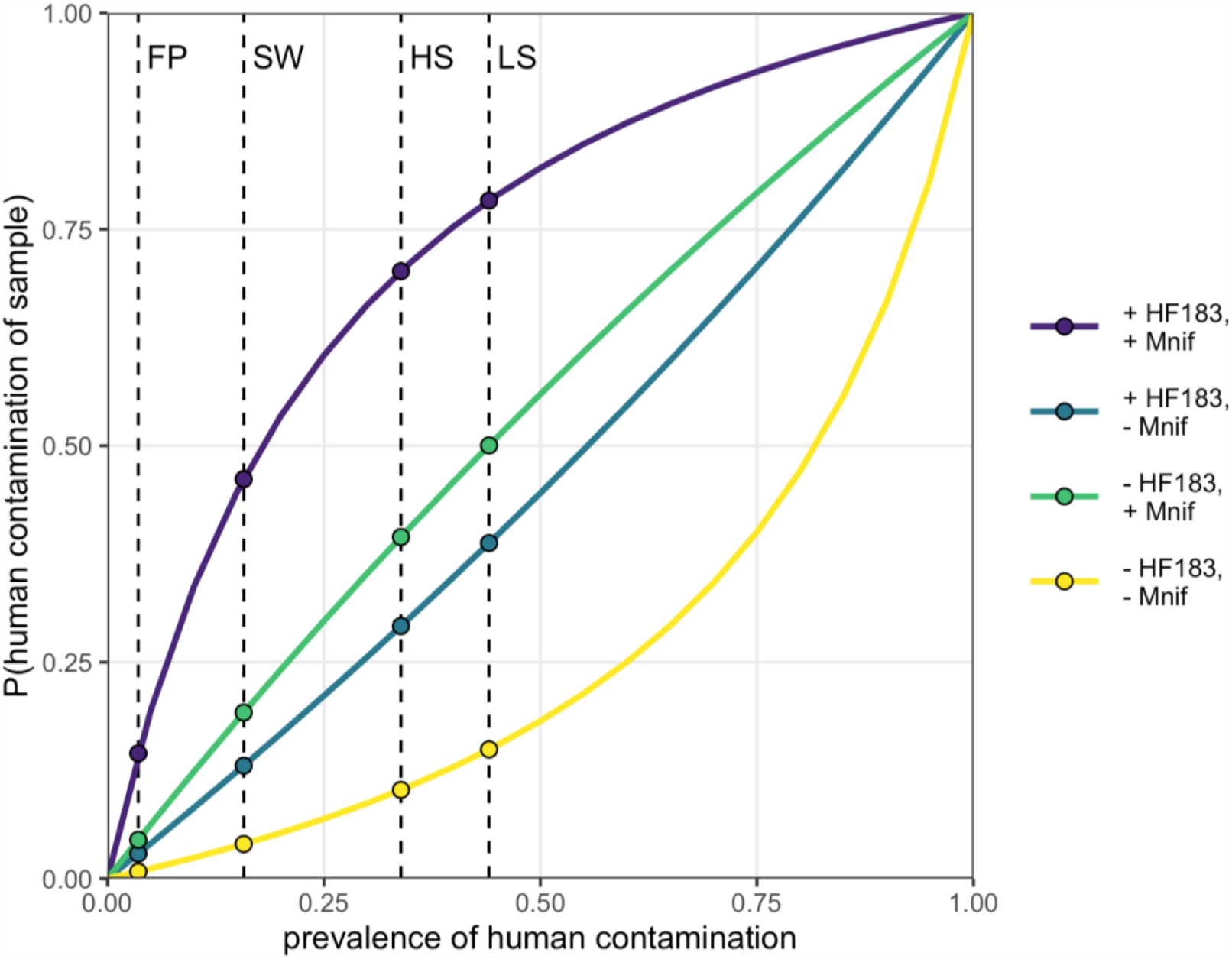
Conditional probability of sample contamination with human feces given detection status of both HF183 and Mnif for all values of human contamination prevalence. Values of sensitivity and specificity were obtained using human and animal feces from the study area, and are 64% and 67%, respectively, for HF183 and 71% and 70% for Mnif. The dashed vertical lines indicate the HF183 detection frequency for each sample type to illustrate relevant human contamination probabilities. FP: food preparation surfaces; SW: stored water; HS: household entrance soil; LS: latrine entrance soil.

### Prevalence of human fecal contamination

Posterior predictions from each of the five accuracy-adjusted models were used to estimate stratum-specific prevalence of human fecal contamination. To compare treatment assignments and study phases, we predicted prevalence for compounds with no animals or antecedent precipitation and the sample mean population density (7 persons/100 m^2^), wealth score (46), and previous-day temperature (20.4 °C), in which soil surfaces were dry and shaded and water storage containers possessed wide, uncovered mouths. The prevalence estimates were notably imprecise; the 95% CI of the HF183 prevalence in post-treatment latrine soil ranged from 3% to 92% for Model 2 (Table 2). The 95% CI widths were similar for Model 1 and the bootstrap estimates but were substantially wider for the other four models, which accounted for FST marker sensitivity and specificity (see SI). The intervals narrowed somewhat when both indicators were considered (Model 4) and narrowed further when all sample types were incorporated (Model 5) but were still wider than the estimates that did not account for diagnostic accuracy.

**Table 2.**
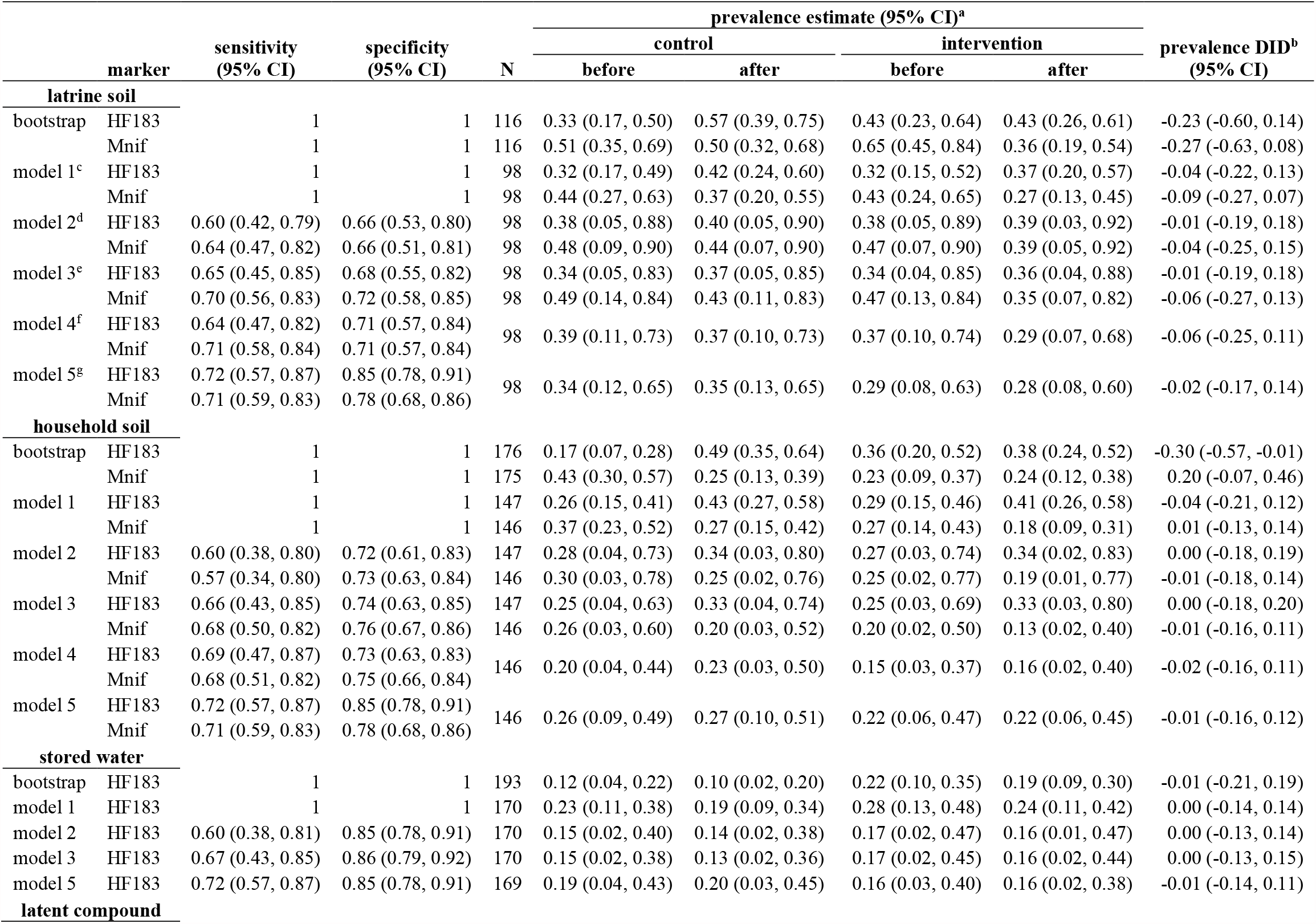
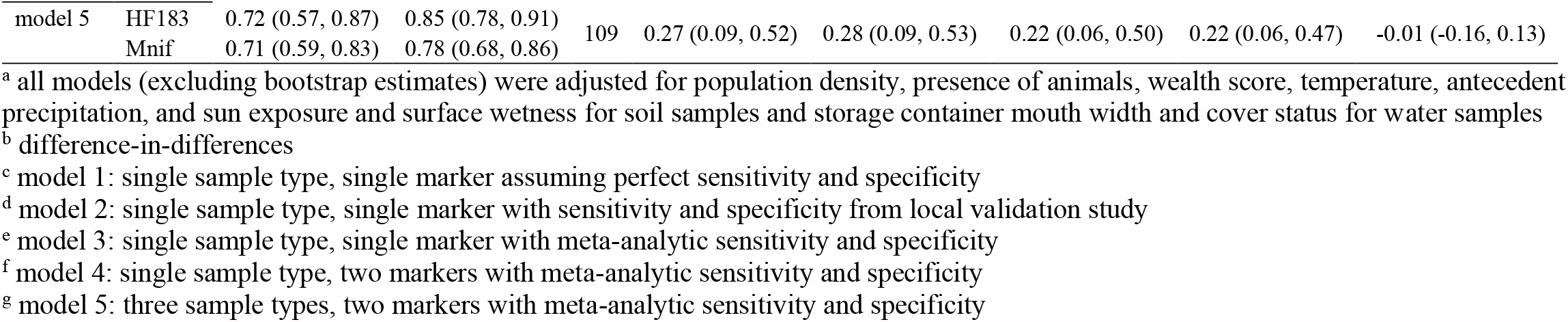
Bootstrap and adjusted model-based estimates human marker sensitivity and specificity, prevalence of human fecal contamination stratified by treatment arm and study phase, and effect of the sanitation intervention on human fecal contamination prevalence in soil and water from MapSan study compounds

Although we did not formally assess the pairwise differences between prevalence estimates, the wide and largely overlapping posterior predictive CIs indicate a limited ability to distinguish between prevalence estimates between different strata or models. The DID estimates on the probability scale were strongly consistent with no effect for all model specifications, which further suggests that the available data were insufficient to assess prevalence differences between strata. The corresponding prevalence odds ratio estimates obtained directly from the DID product term were likewise imprecise (Figure S1). Nonetheless, the model-based prevalence estimates were consistently more similar between study phase and treatment group than the corresponding bootstrap estimates. This trend was notable for Model 5, which assumed that time and treatment effects acted directly on the compound-wide prevalence of human contamination, thus affecting all three sample types equally. The compound-level prevalence estimates were quite similar, particularly between study phases for the same treatment group: 27% (95% CI: 9-52%) at baseline and 28% (9-53%) at follow-up for control compounds and 22% (6-50%) at baseline and 22% (6-47%) at follow-up for intervention compounds. The corresponding estimates for household soil were nearly identical to the compound-level estimates, with somewhat higher estimates for latrine soil and lower for stored water. Although the physical interpretation of this compound-level construct is uncertain, these estimates suggest that about a quarter of compounds were measurably impacted by human fecal contamination, which was unaffected by improvements to shared sanitation facilities.

## Discussion

The provision of shared latrines reduced average soil concentrations of the molecular *E. coli* marker EC23S at latrine entrances by more than 1-log_10_ but did not have a comparable effect on culturable *E. coli*. EC23S latrine soil concentrations rose more in control compounds than they fell in intervention compounds, which under the parallel trends assumption is interpreted as a secular trend upwards that the intervention mitigated, for a much smaller absolute reduction than suggested by the DID estimate (Figure 1).^43^ However, an opposite, downward trend was observed for all cEC concentrations. This discrepancy between two tests for the same organism complicates the interpretation of the relatively strong intervention effect estimated for EC23S. While the exact reasons for this discrepancy are yet to be determined, preliminary evidence from a related analysis suggests that the modified mTEC broth used for *E. coli* culture may have produced colonies of the same color and morphology for *Klebsiella* spp., which are commonly soil-derived and not specific to feces.^70^ By contrast, the developers of EC23S reported 95% specificity to *E. coli* and cross reactions only with other *Escherichia* species, not *Klebsiella*.^46^ Accordingly, EC23S potentially better reflected trends in fecal contamination, while cEC may have been confounded by soil microbes more susceptible to environmental conditions, such as the 2016 drought in southern Mozambique.^71^

A cluster-randomized trial in rural Bangladesh likewise found scant evidence of reductions in culturable *E. coli* concentrations from sanitation improvements.^72,73^ Latrine provision also did not reduce the prevalence of pathogenic *E. coli* genes in soil, meaning neither culture-nor molecular-based measurements of soil *E. coli* were affected.^39^ Other recent trials have not assessed intervention impacts on fecal contamination of soil, but several have evaluated contamination of drinking water, with some also testing child hands, food, or fomites.^15^ As with the present study, all found no effect of sanitation-only interventions on any environmental compartment; combined water, sanitation, and hygiene interventions improved drinking water quality in two studies.^13,14^

Measures of human-associated FST markers demonstrated that about a quarter of compounds were impacted by human fecal contamination, with compound-level prevalence estimates not statistically different at baseline and follow-up. Similarly, two cluster-randomized trials, in India and Bangladesh, found no effect of rural sanitation interventions on the prevalence of human-associated indicators in stored drinking water.^37,39^ Both studies also assessed human markers in mother and child hand rinse samples, which were not collected in this study. No effect was observed for either hand type in India or on mother hands in Bangladesh, although the human marker prevalence may have been reduced on child hands.^39^

Accounting for the diagnostic accuracy of FST markers revealed far greater uncertainty about host-specific fecal contamination, both of individual samples and population averages, than indicated by the raw indicator measurements. The relatively poor sensitivity and specificity of both human markers in this setting severely limited their ability to identify specific samples contaminated with human feces, but even moderate improvements in accuracy could substantially increase FST marker utility. For example, a study in Singapore reported 75% sensitivity and 89% specificity for HF183,^74^ corresponding to a 55% chance a positive sample is contaminated at 15% background prevalence and an 84% chance at 44% prevalence, compared with 26% and 60%, respectively, for detection of HF183 in our study. Correcting for indicator sensitivity and specificity to human-source contamination, coupled with the limited observations of each sample type, yielded imprecise prevalence estimates that were consistent with both near absence and almost omnipresence of contamination. While the reduced amplification efficiency of HF183 (82%) may have contributed to its low sensitivity, it produced similar accuracy-corrected estimates as Mnif, which was 95% efficient (Table S5). This imprecision inhibited detecting intervention effects. The point estimates for the intervention effect were relatively close to the null but the full posterior distributions were consistent with both large reductions and substantial increases in prevalence attributable to the intervention. This analysis does not rule out the possibility that sanitation improvements reduced the prevalence of human fecal contamination. Rather, it strongly suggests that the tools used were inadequate, conveying too little information to address the research question with an acceptable degree of confidence.

These limitations highlight the importance of conducting local validation studies for any new FST application.^75^ Accounting for diagnostic accuracy is unlikely to improve the strength or precision of estimates, but may help mitigate overconfidence and overinterpretation by revealing limitations of the available measurements. This practice could also be extended to account for indicator sensitivity and specificity to strictly fecal targets, rather than environmental microbes with non-fecal origins, although we lacked the appropriate data to implement such an analysis for our two non-specific indicators, EC23S and cEC. As the diagnostic accuracy framework is currently limited to binary outcomes, analysis of such high-prevalence indicators would benefit from the development of analogous approaches for continuous outcomes. Given the intermingling in low-income settings of humans and animals, and their gut microbiomes, alternative FST targets such as mitochondrial DNA could prove more accurate.^76,77^ Recent technological advances also present opportunities for new approaches that might bypass the limitations of the current FST paradigm, including portable, long-read sequencing platforms for metagenomic-based source tracking and parallel PCR platforms that render simultaneous analysis of multiple FST markers and comprehensive direct pathogen detection increasingly feasible.^20,78–82^ These technologies will also need to overcome the substantial variability, limited analytical sensitivity, and matrix interference characteristic of environmental microbial assessments.^16^

The low signal typical of environmental measurements suggests that study designs— preferably longitudinal—that maximize observations on select pathways of greatest interest should be prioritized to support more robust inference, regardless of analytical approach.^83^ A recent longitudinal analysis of *E. coli* concentrations in rural Bangladesh, collected at eight timepoints over 2.5 years from 720 households, demonstrates the advantages of maximizing the number of basic measurements across time. Although pooled estimates from certain sample types achieved statistical significance, the sheer quantity of information available convincingly demonstrated the lack of a physically meaningful sanitation intervention impacts on ambient fecal contamination.^73^

Many have speculated that sanitation’s apparent lack of effect may be due in part to animal fecal contamination.^12,22^ Animal feces often contain pathogens capable of infecting humans and animal fecal biomass in domestic environments is estimated to far exceed that from humans. ^22,84–86^ Inadequate management of child feces and fecal sludge, contamination of food and water outside the home, and inadequate community-level drainage, solid waste, and sanitation services all present potential pathways of continued contamination despite household sanitation improvements.^24,87–92^ Recognizing calls for “transformative” WASH to address these multifarious hazards, sustained progress may require high standards of housing and public services in addition to WASH improvements, necessitating multi-sectoral coordination and financing.^12,93–95^ Even small treatment effects may translate to positive economic benefits.^12^ Additionally, quality sanitation infrastructure can provide important benefits irrespective of preventing pathogen exposure, particularly in crowded urban settlements.^96,97^ For example, previous research found users of MapSan intervention latrines and similar facilities in the same neighborhoods reported reduced disgust and embarrassment about unhygienic conditions and improved perceptions of security and privacy.^98^ Based on the results of our study, we recommend future research to understand the etiology and ecology of fecal pathogens in domestic environments and beyond to help inform interventions needed to construct healthy environments and to protect children’s health.

## Acknowledgment

This study was funded by the United States Agency for International Development under Translating Research into Action, Cooperative Agreement No. GHS-A-00-09-00015-00, the Bill and Melinda Gates Foundation grant OPP1137224, and the National Institute of Environmental Health Sciences (T32ES007018). We gratefully acknowledge our implementing partner, Water and Sanitation for the Urban Poor; the field research team at WE Consult; the laboratory staff at the Mozambique National Institute of Health; and Olimpio Zavale, Olivia Ginn, and Anna Stamatogiannakis for sampling support.

## Supporting Information Available

Site selection criteria; samples collected; qPCR assay details; calibration curves; detection limits; laboratory quality control; conditional probability; difference-in-differences estimates; validation studies; diagnostic accuracy; accuracy-adjusted intervention effect estimates; human fecal contamination prevalence estimates; prior distributions; model Stan code.

