## Supporting Information for "Impacts of an urban sanitation intervention on fecal indicators and the prevalence of human fecal contamination in Mozambique"

This supporting information contains 47 pages, 1 figure, and 10 tables.

### S1. Site selection criteria

Intervention sites were selected by the nongovernmental organization (NGO) Water and Sanitation for the Urban Poor (WSUP) according to feasibility and demand criteria (Table S1). The MapSan researchers were not involved in the design of the intervention or selection of the intervention sites, but did recruit similar compounds to serve as control sites according to a reduced set of the same selection criteria applied to intervention sites.

**Table S1. Baseline site selection criteria for intervention and control compounds**

| | required for | |
|---|---|---|
| **criterion** | **intervention compounds** | **control compounds** |
| located in the 11 pre-defined implementation neighborhoods | yes | no[a] |
| residents share sanitation in poor condition | yes | yes |
| at least 12 residents | yes | yes |
| residents willing to contribute to latrine construction costs | yes | yes |
| sufficient space available for construction of the new facility | yes | no |
| accessible for transportation of construction materials and tank-emptying activities | yes | no |
| legal access to piped water supply | yes | yes |
| groundwater level deep enough to accommodate a septic tank | yes | no |
| at least one child younger than 48 months old in residence | no | yes |

[a] also recruited from 6 similar, adjacent neighborhoods; see Knee et al. (2021)[1]

All children under 48 months old living in study compounds at baseline enrollment were invited to participate in the MapSan health impact trial following written, informed consent from a parent or guardian. At 12-month follow-up, any children living in study compounds who were not previously enrolled but would have been under 48 months of age at the time the compound was enrolled at baseline were also invited to participate, including children born after baseline. Knee et al. (2021) provide additional details on eligibility and enrollment into the MapSan child health study component.[1]

We collected environmental samples from a subset of compounds already scheduled for baseline enrollment during May – August 2015. The specific intervention compounds at which we collected baseline environmental samples were selected opportunistically, largely prioritizing compounds with visits scheduled earlier in the morning to ensure sufficient time for sample transport and laboratory processing. Control compounds were similarly selected for environmental sampling among those already scheduled for baseline enrollment, although we prioritized control compounds with similar numbers of residents as the intervention compounds that had been selected for environmental sampling in the preceding two weeks.

The compounds selected for environmental sampling at baseline were revisited in June – September 2016, 12 months (± 2 weeks) following the opening of the new latrine for intervention compounds and 12 months (± 2 weeks) after baseline enrollment for control compounds. Four compounds at which environmental samples were collected at baseline were unavailable at follow up due to travel or relocation of eligible children for the health impact study. The provision of intervention latrines was substantially delayed for two additional compounds following baseline enrollment, rendering them outside the 12-month (± 2 weeks) follow-up window for the duration of the June – September 2016 environmental sampling campaign. However, environmental samples were collected from 13 additional compounds during the 12-month follow-up phase that had not been sampled at baseline but had been enrolled in the larger MapSan trial at baseline. These additional compounds were selected opportunistically in the same manner as the initial baseline sample set, but intervention compounds (n = 11) were prioritized over control compounds (n = 2), which had been overrepresented at baseline.

**S2. Samples collected**

We visited additional households and compounds in the follow-up study phase, collecting more samples than at baseline (Table S2).[2] Fewer source water samples were collected at follow-up because the municipal water supply was available less often. One control compound (with two enrolled households) independently upgraded the latrine after baseline and was excluded from the follow-up sample.

**Table S2. Number of observations of compounds, households, and each sample type**

| | before | | after | |
|---|---|---|---|---|
| **observation unit** | **control** | **intervention** | **control** | **intervention** |
| compounds | 33 | 25 | 30 | 34 |
| households | 86 | 66 | 86 | 90 |
| latrine entrance soil | 33 | 23 | 30 | 30 |
| household entrance soil | 49 | 36 | 47 | 45 |
| compound source water | 23 | 21 | 19 | 22 |
| household stored water | 50 | 41 | 52 | 55 |
| food preparation surfaces | 50 | 40 | 53 | 51 |

We requested that respondents provide both household stored water and household food preparation surfaces in the manner or condition in which they would typically be used. For stored water, we requested that the respondent provide it to us as if they were giving a child water to drink. Similarly, we requested that the household respondent identify or provide a surface they would typically use to prepare food, in the condition in which they would use it. If multiple surfaces were available, the respondent decided which to present as representative of their ordinary food preparation practices. Additional descriptions of specific sampling procedures and the observed baseline characteristics of each sample type are provided in Holcomb et al. (2020).[2]

#### S3. qPCR assay details

All qPCR assays were performed using TaqMan Environmental Master Mix 2.0 and were subjected to an initial 10 minute incubation at 95°C, after which the cycling conditions specified by the assay developers were followed for each assay (Table S3).

**Table S3. qPCR assay details**

| assay[reference] | cycles | parameters | oligonucleotide (nM) | sequence (5'-3') |
|---|---|---|---|---|
| EC23S857[3] | 40 | 15 s: 95 °C<br>60 s: 60 °C | F (1000) | GGTAGAGCACTGTTTtGGCA |
| | | | R (1000) | TGTCTCCCGTGATAACtTTCTC |
| | | | P (80) | 6-FAM-TCATCCCGACTTACCAACCCG-BHQ1 |
| HF183/ BacR287[4] | 40 | 15 s: 95 °C<br>60 s: 60 °C | HF183 (1000) | ATCATGAGTTCACATGTCCG |
| | | | BacR287 (1000) | CTTCCTCTCAGAACCCCTATCC |
| | | | BacP234MGB (80) | 6-FAM-CTAATGGAACGCATCCC-BHQplus |
| Mnif [5] | 50 | 10 s: 95 °C<br>30 s: 57 °C | Mnif-202F (800) | GAAAGCGGAGGTCCTGAA |
| | | | Mnif-353R (800) | ACTGAAAAACCTCCGCAAAC |
| | | | Mnif-236P (240) | 6-FAM-CCGGACGTGGTGTAAC AGTAGCTA-BHQ1 |
| Sketa22[6,7] | 40 | 15 s: 95 °C<br>60 s: 60 °C | SketaF2 (1000) | GGTTTCCGCAGCTGGG |
| | | | SketaR22 (1000) | CCGAGCCGTCCTGGTC |
| | | | SketaP2 (80) | 6-FAM-AGTCGCAGGCGGCCACCGT-BHQ1 |

### S4. Calibration curves

Calibration curves for quantifying molecular fecal indicator gene copies were fit to ten-fold dilution series of positive controls (PC) that had been spiked with reference material for all three assays and extracted alongside each batch of samples.[2] Three gBlock linear DNA fragments (Integrated DNA Technologies, Skokie, Il, USA) containing composite reference sequences for the three fecal source tracking assays considered in this paper as well as additional assays considered in the associated validation study were used as standard reference material for the positive controls (Table S4).[2,8,9] Extracting the reference material accounted for loss of target DNA during extraction but induced additional variability between dilution series constructed from each PC. We therefore allowed both the slopes and intercepts of the calibration curves to vary by qPCR instrument run and extraction batch to account for the additional variation. The resulting curves were relatively linear ($R^2$ > 95%) when averaged across all instrument runs and extraction batches but were somewhat inefficient, particularly for HF183 (Table S5).

**Table S4. Synthetic DNA reference material spiked into positive controls**

| assays covered | sequence [5'-3'] | GenBank (base positions) | length [bases] |
|---|---|---|---|
| BacHum-UCD[10]<br>BacUni-UCD[10]<br>HF183/BacR287[11] | CCAGGATGGGATCATGAGTTCACATGTCCGCATGAT<br>TAAAGGTATTTTCCGGTAGACGATGGGGATGCGTTC<br>CATTAGATAGTAGGCGGGGTAACGGCCCACCTAGTC<br>AACGATGGATAGGGGTTCTGAGAGGAAGGTCCCCCA<br>CATTGGAACTGAGACACGGTCCAAACTCCTACGGGA<br>GGCAGCAGTGAGGAATATTGGTCAATGGGCGATGGC<br>CTGAACCAGCCAAGTAGCGTGAAGGATGACTGCCCT<br>ATGGGTTGTAAACTTCTTTTATAAAGGAATAAAGTC<br>GGGTATGCATACCCGTTTGCATGTACTTTATGAATAA<br>GGATCGGCTAACTCCGTGCCAGCAGCCGCGGTAATA<br>CGGAGGATCCGAGCGTTATCCGGATTTATTGGGTTT<br>AAAGGGAGCGTAGATGGATGTTTAAGTCAGTTGTGA<br>AAGTTTGCGGCTCAACCGTAAAATTGCAGTTGATAC<br>TGGATGTCTTGAGTGCAGTTGAGGCAGGCGGAATTC<br>GTGGTGTAGCGGTGAAATGCTTAGATATCACGAAGA<br>ACTCCGATTGCGAAGGCAGC | AB242142<br>(170-730) | 560 |
| GFD[12]<br>LA35[13] | TGGGTCTAATACCGGATACGACCATCTGCCGCATGG<br>CGGGTGGTGGAAAGTTTTTCGATTGGGGATGGGCTC<br>GCGGCCTATCAGTTTGTTGGTGGGGTAATGGCCTAC<br>CAAGGCGACGACGGGTAGCCGGCCTGAGAGGGCGA<br>CCGGCCACACTGGGACTGAGACACGGCCCAGACTCC<br>TACGGGAGGCAGCAGTGGGGAATATTGCACAATGG<br>GGGAAACCCTGATGCAGCGACGCAGCGTGCGGGAT<br>GACGGCCTTCGGGTTGTAAACCGCTTTCAGCAGGGA<br>AGAAGCCTTCGGGTGACGGTACCTGCAGAAGAAGTA<br>CCGGCTAACTACGTGCCAGCAGCCGCGGTAATACGT<br>AGGGTACGAGCGTTGTCCGGAATTATTGGGCGTAAA<br>GAGCTCGTAGGTGGTTGGTCACGTCTGCTGTGGAAA<br>CGCAACGCTTAACGTTGCGCGGGCAGTGGGTACGGG<br>CTGACTAGAGTGCAGTAGGGGAGTCTGGAATTCCTG<br>GTGTAGCGGTGAAATGCGCAGATATCAGGAGGAAC<br>ACCGGTGGCGAAGGCGGGACTCTGGGCTGTGACTGA<br>CACTGGGGAGCGAAAGCATTGCTAACAGTTcGGCTG<br>AGCACTCTAGGGAGACTGCCTTCGCAAGGAGGAGGA<br>AGGTGAGGACGACGTCAAGTCATCATGGCCCTTACG<br>CCTAGGGCTACACACGTGCTACAATGGGATGTACAA<br>AGAGACGCAATACCGCGA | FJ462358<br>(156-746)<br>JN084061<br>(29-171) | 732 |
| EC23S857[3]<br>HAdV[14]<br>Mnif[5] | TAACTATGGTCATCGTTCGTCAGCAGTAACAGTAATT<br>GCTACACCTGCTGAAACCACTGTCCCTTTTTCTTGGG<br>CAACTCTTGTTTATGTGTTGAAAGCGGAGGTCCTGA<br>ACCGGGTGTTGGCTGTGCCGGACGTGGTGTAACAGT<br>AGCTATGAAAAGACTTGAAAACTTAGGTGTTTTTGA<br>TAAGGATTTGGATGTAGTCATTTATGGTGTACTTGGA<br>GATGTTGTTTGCGGAGGTTTTTCAGTGCCTTTACGTT<br>CTCGGGCCAGGACGCCTCGGAGTACCTGAGCCCCGG<br>GCTGGTGCAGTTTGCCCGCGCCACCGAGACGTACTT<br>CAGCCTGAATAACAAGTTTAGAAACCCCACGGTGGC<br>GCCTACGCATCTCCGGGGGTAGAGCACTGTTTCGGC<br>AAGGGGGTCATCCCGACTTACCAACCCGATGCAAAC<br>TGCGAATACCGGAGAATGTTATCACGGGAGACACAC<br>GGCGGGT | AE015928<br>(4515891-<br>4515973)<br>AB019138<br>(192-363)<br>AC_000008<br>(18885-19000)<br>DQ682619<br>(847-954) | 475 |

**Table S5. Mean (95% CI) estimates of calibration curve parameters**

| assay | intercept | slope | efficiency (%) | $R^2$ |
|---|---|---|---|---|
| EC23S | 47.9 (47.3, 48.6) | -3.50 (-3.64, -3.37) | 93.1 (88.1, 98.1) | 0.98 (0.97, 0.98) |
| HF183 | 47.5 (46.4, 48.6) | -3.85 (-4.07, -3.67) | 81.8 (76.1, 87.4) | 0.98 (0.97, 0.98) |
| Mnif | 48.8 (47.7, 50.3) | -3.47 (-3.79, -3.23) | 94.5 (83.6, 104.0) | 0.95 (0.93, 0.95) |

### S5. Detection limits

The limit of detection (LOD) for each assay was obtained in terms of the quantification cycle (Cq), the number of amplification cycles above which the target would be considered absent from the reaction, using receiver operating characteristic (ROC) analysis.[2,15] We performed ROC analysis on the local validation study data presented in Holcomb et al. (2020), considering Cq cutoff values from 10 to the maximum number of cycles indicated by the assay developers, in full-cycle increments. We calculated diagnostic sensitivity and specificity for each cutoff value, considering any reactions with a Cq value below the cutoff as positive. The highest Cq value that maximized the Youden index, $J = sensitivity + specificity - 1$, for each assay was considered the assay LOD (Table S6).[15,16]

Process limits of detection (PLOD) were estimated for each sample from the assay LOD Cq values, the extraction batch- and instrument run-specific calibration curve estimates, and the amount of sample processed, i.e. volume of water filtered or the dry mass or surface area represented by the eluate filtered for soil samples and surface swabs, respectively. For *E. coli* enumerated by culture (cEC), we assumed an assay LOD of 1 cfu per plate and calculated the PLOD for each sample on the basis of the amount of sample represented by the least-diluted plate read for that sample.[2] The PLOD averages indicate that relatively high gene copy concentrations were required for reliable detection by any of the three qPCR-based assays, on the order of >10,000 gc/dry gram of soil, nearly 1000 gc/100 mL water, and >3000 gc/100 $cm^2$ of food preparation surface (Table S6).

**Table S6. qPCR assay limits of detection and mean sample-specific process limits of detection by sample matrix**

| | | process limit of detection | | | | | |
|---|---|---|---|---|---|---|---|
| | | soil | | water | | surface | |
| | assay | $log_{10}$ gc[c]/dry g | | $log_{10}$ gc/100 mL | | $log_{10}$ gc/100 $cm^2$ | |
| **assay** | **LOD[a] Cq[b]** | **n** | **mean (SD[d])** | **n** | **mean (SD)** | **n** | **mean (SD)** |
| EC23S | 39 | 298 | 4.51 (0.11) | 292 | 3.27 (0.12) | 199 | 3.96 (0.30) |
| HF183 | 39 | 299 | 4.24 (0.33) | 292 | 2.85 (0.29) | 199 | 3.45 (0.36) |
| Mnif | 41 | 298 | 4.18 (0.17) | 291 | 3.03 (0.11) | 196 | 3.53 (0.35) |

[a] limit of detection
[b] quantification cycle
[c] gene copies
[d] standard deviation

### S6. Laboratory quality control

Sterile PBS was filtered as a laboratory blank for approximately every 10 samples filtered, all of which were culture-negative for cEC (n = 151). At least three no template control (NTC) reactions were included on every qPCR run using 5 µL nuclease-free water in place of sample template. Each qPCR run typically included samples from three extraction batches, and a negative extraction control (NEC) was included from each extraction batch represented on the plate. HF183 was absent in all NTC (n = 46) and NEC (n = 46) reactions. Mnif was likewise not detected in any NTC (n = 42) or NEC (n = 46) reaction. However, EC23S was detected in 2% of NTC (1/45) and 11% of NEC (5/46) reactions. EC23S concentrations were low in the contaminated negative controls, with a mean Cq of 38.1 and a minimum Cq of 37.1—only slightly above the detection limit of 39 cycles. Such low levels of contamination have been reported previously and are thought to be due to residual *E. coli* DNA in the Environmental Master Mix from the production process.[2,17] One latrine soil sample was considered inhibited based on a greater than 3 Cq deviation from the mean Sketa22 Cq of all NECs and positive controls on the same plate. This sample was diluted 1:5 in all further analyses. For each sample type, 10% of samples were randomly selected to be analyzed by each qPCR assay in technical duplicate reactions. The detection status of EC23S agreed in 96% (74/77) of replicate pairs.

HF183 results matched for 86% (66/77) of paired reactions and Mnif agreed in 95% (71/75). Samples analyzed in duplicate were considered positive for a given target if at least one of the replicate reactions was above the limit of detection.

**S7. Conditional probability**

The conditional probability of human contamination, $C$, given the detection of two human-associated markers, $M_1$ and $M_2$ is given by

$$
\begin{aligned}
P(C^+|M_1^+ \cap M_2^+) &= \frac{P(M_1^+|C^+) \times P(M_2^+|C^+) \times P(C^+)}{P(M_1^+|C^+) \times P(M_2^+|C^+) \times P(C) + P(M_1^+|C^-) \times P(M_2^+|C^-) \times P(C^-)} \\
&= \frac{Se_1 \times Se_2 \times P(C^+)}{Se_1 \times Se_2 \times P(C^+) + (1 - Sp_1) \times (1 - Sp_2) \times \left(1 - P(C^+)\right)}
\end{aligned} \quad \textbf{(1)}
$$

where $Se_1 = P(M_1^+|C^+)$, $Se_2 = P(M_2^+|C^+)$, $Sp_1 = P(M_1^-|C^-)$, $Sp_2 = P(M_2^-|C^-)$, and $A^+$ indicates the presence, and $A^-$ the absence, of variable $A$.[18] We calculated the conditional probability of contamination for HF183 and Mnif separately and for each combination of the two indicators ($M_1^+, M_2^+$; $M_1^+, M_2^-$; $M_1^-, M_2^+$; $M_1^-, M_2^-$) by sample type.

**S8. Difference-in-differences estimates**

The effect of the intervention was estimated using difference-in-differences approaches. Crude DID estimates of *E. coli* $\log_{10}$-concentration and human FST marker prevalence were calculated using 2000 bootstrap samples (Table S7). The DID estimate was calculated as

$$DID = \left(\mathrm{E}\left[Y_{T=1,P=1}\right] - \mathrm{E}\left[Y_{T=1,P=0}\right]\right) - \left(\mathrm{E}\left[Y_{T=0,P=1}\right] - \mathrm{E}\left[Y_{T=0,P=0}\right]\right) \quad \textbf{(2)}$$

where $Y_{T,P}$ are the observed indicator values for treatment group $T$ in study phase $P$. The value of $T$ is 0 for control compound observations and 1 for intervention compounds; likewise, $P$ takes the value 0 for pre-treatment (baseline) observations and 1 post-treatment (follow-up). Observations below the PLOD or too numerous to count were imputed by sample type from a truncated normal distribution with mean and standard deviation obtained through maximum likelihood estimation, assuming the $\log_{10}$ concentration was normally distributed and subject to left- and right-censoring.[2,19] Model-based DID estimates were obtained using the product-term representation of the DID estimator in regression models, which permits the inclusion of additional covariates. We produced both crude model-based estimates, which only included terms for the DID estimator, as well as adjusted estimates that included additional terms for meteorological, compound, household, sample characteristics (Table S8).

**Table S7. Bootstrap difference-in-differences estimates**

| | | before | | | | after | | | | |
|---|---|---|---|---|---|---|---|---|---|---|
| | | control | | intervention | | control | | intervention | | DID[a] |
| indicator | sample type | N | estimate | N | estimate | N | estimate | N | estimate | estimate |
| ***E. coli*** **$log_{10}$ concentration, mean (95% CI)** | | | | | | | | | | |
| cEC | latrine soil | 33 | 4.0 (3.6, 4.3) | 23 | 4.0 (3.5, 4.4) | 30 | 3.3 (2.9, 3.7) | 30 | 3.0 (2.6, 3.5) | -0.3 (-1.1, 0.5) |
| | household soil | 49 | 4.1 (3.8, 4.3) | 36 | 4.2 (3.9, 4.5) | 47 | 3.7 (3.5, 4.0) | 45 | 3.4 (3.1, 3.8) | -0.4 (-1.0, 0.1) |
| | stored water | 50 | 1.8 (1.4, 2.2) | 41 | 1.6 (1.1, 2.0) | 52 | 1.1 (0.7, 1.4) | 55 | 1.0 (0.7, 1.3) | 0.2 (-0.6, 0.9) |
| | food surface | 50 | 3.3 (2.8, 3.8) | 40 | 2.9 (2.3, 3.6) | 53 | 2.0 (1.5, 2.5) | 51 | 1.9 (1.4, 2.4) | 0.3 (-0.8, 1.3) |
| EC23S | latrine soil | 33 | 6.2 (5.8, 6.6) | 23 | 6.9 (6.4, 7.4) | 30 | 6.7 (6.4, 7.1) | 30 | 6.5 (6.0, 7.0) | -0.9 (-1.8, 0.0) |
| | household soil | 49 | 6.8 (6.5, 7.0) | 36 | 6.7 (6.4, 7.0) | 47 | 6.7 (6.4, 7.0) | 45 | 6.6 (6.4, 6.9) | 0.0 (-0.5, 0.5) |
| | stored water | 50 | 4.1 (3.9, 4.3) | 41 | 4.4 (4.2, 4.7) | 52 | 4.2 (3.9, 4.4) | 55 | 4.0 (3.8, 4.2) | -0.4 (-0.9, 0.0) |
| | food surface | 50 | 4.4 (4.2, 4.7) | 40 | 5.0 (4.8, 5.3) | 53 | 4.7 (4.4, 5.0) | 51 | 4.5 (4.2, 4.8) | -0.8 (-1.4, -0.3) |
| **human marker prevalence, % (95% CI)** | | | | | | | | | | |
| HF183 | latrine soil | 33 | 33 (17, 50) | 23 | 43 (23, 64) | 30 | 57 (39, 75) | 30 | 43 (26, 61) | -23 (-60, 14) |
| | household soil | 49 | 17 (7, 28) | 36 | 36 (20, 52) | 47 | 49 (35, 64) | 45 | 38 (24, 52) | -30 (-57, -1) |
| | stored water | 50 | 12 (4, 22) | 41 | 22 (10, 35) | 52 | 10 (2, 20) | 55 | 19 (9, 30) | -1 (-21, 19) |
| | food surface | 50 | 2 (0, 7) | 40 | 0 (0, 0) | 53 | 9 (2, 18) | 51 | 2 (0, 7) | -5 (-16, 4) |
| Mnif | latrine soil | 33 | 51 (35, 69) | 23 | 65 (45, 84) | 30 | 50 (32, 68) | 30 | 36 (19, 54) | -27 (-63, 8) |
| | household soil | 49 | 43 (30, 57) | 36 | 23 (9, 37) | 47 | 25 (13, 39) | 45 | 24 (12, 38) | 20 (-7, 46) |
| | stored water | 50 | 0 (0, 0) | 41 | 2 (0, 8) | 52 | 0 (0, 0) | 55 | 0 (0, 0) | -2 (-8, 0) |
| | food surface | 50 | 0 (0, 0) | 40 | 3 (0, 8) | 53 | 2 (0, 7) | 51 | 0 (0, 0) | -4 (-12, 0) |

[a] difference-in-differences: (intervention, after – intervention, before) – (control, after – control, before)

**Table S8. Model-based difference-in-differences estimates**

| | latrine entrance soil | | | | household entrance soil | | | | household stored water | | | | food preparation surfaces | | | |
|---|---|---|---|---|---|---|---|---|---|---|---|---|---|---|---|---|
| | crude[a] | | adjusted[b] | | crude | | adjusted | | crude | | adjusted | | crude | | adjusted | |
| **target** | **N** | **DID[c]** | **N** | **DID** | **N** | **DID** | **N** | **DID** | **N** | **DID** | **N** | **DID** | **N** | **DID** | **N** | **DID** |
| ***E. coli*** **$\log_{10}$ concentration change (95% CI)** | | | | | | | | | | | | | | | | |
| cEC | 111 | -0.37 (-1.17, 0.44) | 95 | -0.42 (-1.28, 0.40) | 175 | -0.34 (-0.92, 0.22) | 146 | 0.05 (-0.62, 0.72) | 194 | 0.15 (-0.61, 0.92) | 170 | -0.42 (-1.28, 0.44) | 192 | 0.18 (-0.9, 1.26) | 169 | -0.11 (-1.4, 1.17) |
| EC23S | 116 | -0.84 (-1.64, -0.02) | 98 | -1.22 (-2.11, -0.30) | 176 | 0.06 (-0.46, 0.57) | 147 | 0.36 (-0.26, 1.01) | 193 | -0.46 (-0.89, -0.04) | 170 | -0.41 (-0.92, 0.12) | 193 | -0.82 (-1.46, -0.19) | 171 | -0.56 (-1.32, 0.19) |
| **human target prevalence odds ratio (95% CI)** | | | | | | | | | | | | | | | | |
| HF183 | 116 | 0.94 (0.40, 1.87) | 98 | 0.92 (0.38, 1.87) | 176 | 0.83 (0.39, 1.54) | 147 | 0.90 (0.38, 1.80) | 193 | 1.12 (0.48, 2.27) | 170 | 1.05 (0.44, 2.15) | | | | |
| Mnif | 116 | 0.73 (0.31, 1.44) | 98 | 0.71 (0.29, 1.47) | 175 | 1.24 (0.53, 2.46) | 146 | 1.00 (0.39, 2.06) | | | | | | | | |

[a] not adjusted for covariates; include terms for study phase, treatment arm, and their product, and compound-varying intercepts

[b] adjusted for animal presence, population density, household wealth, temperature, precipitation, and sample-specific variables: sun exposure and surface wetness for soils, presence of lid and width of container mouth for stored water, and food surface material.

[c] difference-in-differences estimated as the regression coefficient on the product of study phase and treatment arm indicators

**S9. Validation studies**

Local differences in diet, geography, and population history affecting the gut microbiome composition of a given population are expected to play the key role in determining local fecal source tracking performance.[20,21] However, multi-laboratory comparisons have also demonstrated that assay performance can vary meaningfully between labs analyzing the same set of challenge samples.[22] Furthermore, assay design and other intrinsic characteristics may also impact potential performance—that is, some assays may have more robust designs that increase the likelihood of performing well in a variety of settings and populations. Source tracking validation studies typically report crude sensitivity and specificity values without quantifying the uncertainty in these estimates, which could be substantial considering the limited number of samples analyzed in many studies, particularly in resource-limited settings (Table S9).

We incorporate a meta-analysis of FST validation studies for our selected human markers, which provides an opportunity to partially pool information across a variety of locations to potentially refine the sensitivity and specificity estimates for our specific study setting.[23] The hierarchical structure of the meta-analytic model means that the estimates for any given location are driven by the data from that location. Sensitivity and specificity estimates at a location with a large amount of data would be almost entirely determined by the local data and largely unaffected by the other studies, while estimates from locations with sparse data, which would be uncertain on their own, incorporate more information pooled from other studies.[24] The extent of information pooling is also determined by the consistency of marker performance across studies. Highly variable marker performance between study locations suggests that performance is mostly driven by differences in local characteristics, limiting the amount of information that can be gained by considering performance other locations. Accordingly, meta-analytic performance estimates in a particular location will be determined largely by the local samples and minimally influenced by data from other locations. On the other hand, similar marker performance across locations suggests that intrinsic assay characteristics play a larger role in its performance, enabling information from other locations to refine the estimates at specific locations with limited local data.[25] We adjust for diagnostic accuracy using the meta-analytic sensitivity and specificity estimates for our study location, such that the estimates are driven by our local data but are influenced by the broader trends in assay performance across all the studies considered. The degree of influence by outside data depends on the degree of uncertainty in the local data (a function of sample size) and the similarity of performance estimates across all the studies considered, which is reflected in the between-study standard deviations, $\sigma^{Se}$ and $\sigma^{Sp}$.

Validation studies included in the performance meta-analysis were identified from Google Scholar records of articles citing the original assay publications and from the references of each published validation study identified.[5,7,11] We only included HF183 studies that assessed the HF183/BFDRev assay or its modification used in this study, HF183/BacR287.[11] We identified HF183 validation studies conducted on five continents, including two in Africa, seven in Asia, two in Australia, four in North America, and one in South America (Table S9). All identified Mnif studies used samples from the USA except for our study in Mozambique. The rural/urban status of the settings from which samples were collected could not be determined for all studies and were more often specified for studies conducted in low- and middle-income countries. We used estimates that considered samples detected below the limit of quantification (DNQ) as negatives when separate estimates were reported by DNQ definition.[26]

**Table S9. Published validation studies of human fecal indicators HF183 and Mnif**

| study | location | setting | human samples N | human samples positive | non-human samples N | non-human samples positive | sensitivity | specificity | ref |
|---|---|---|---|---|---|---|---|---|---|
| **HF183** | | | | | | | | | |
| 1 | Mozambique | urban | 14 | 9 | 27 | 9 | 64% | 67% | [2] |
| 2 | Bangladesh | urban | 5 | 3 | 20 | 12 | 60% | 40% | [21] |
| 3 | Bangladesh | rural | 5 | 4 | 20 | 10 | 80% | 50% | [27] |
| 4 | India | rural | 35 | 10[a] | 60 | 12[a] | 29% | 80% | [28] |
| 5 | Kenya | rural | 17 | 11 | 25 | 0 | 65% | 100% | [29] |
| 6 | Thailand | | 28 | 25 | 100 | 19 | 89% | 81% | [30] |
| 7 | Costa Rica | | 8 | 8 | 47 | 3 | 100% | 94% | [31] |
| 8 | Singapore | urban | 56 | 42 | 85 | 9 | 75% | 89% | [32] |
| 9 | Nepal | | 10 | 10 | 44 | 30 | 100% | 32% | [33] |
| 10 | USA | rural | 4 | 4 | 109 | 0 | 100% | 100% | [34] |
| 11 | Japan | | 20 | 20 | 15 | 0 | 100% | 100% | [35] |
| 12 | Australia | | 32 | 24 | 359 | 12 | 72% | 96% | [36] |
| 13 | USA, Australia | | | | 184 | 6 | | 97% | [37] |
| 14 | USA | | 60 | 57[a] | 130 | 10 [a] | 95% | 92% | [26] |
| 15 | Peru | peri-urban | 30 | 23 | 71 | 24 | 77% | 66% | [38] |
| **Mnif** | | | | | | | | | |
| 1 | Mozambique | urban | 14 | 10 | 27 | 8 | 71% | 70% | [2] |
| 2 | USA | | 60 | 36[a] | 130 | 31[a] | 60% | 76% | [26] |
| 3 | Indiana | | 59 | 40 | 120 | 9 | 68% | 93% | [18] |
| 4 | Mississippi | | 62 | 51 | 243 | 101 | 82% | 58% | [18] |

[a] samples below the limit of quantification considered negative

### S10. Diagnostic accuracy

We considered HF183 validation data from 14 studies of diagnostic sensitivity and 15 studies of specificity (Table S7). For Mnif, we incorporated data from four validation studies. The number of samples ranged from 5–62 for human feces and 15–359 for non-human. Reported crude sensitivity ranged from 29-100% for HF183 and 60-82% for Mnif. Crude specificities were reported from 32–100% for HF183 and 58-93% for Mnif.

By incorporating indicator validation data into models of human fecal contamination, we obtained estimates of indicator sensitivity and specificity that were partially informed by observations in environmental samples with unknown fecal contamination status. For this reason, accuracy estimates from the same model differed by sample type (main text, Table 2). We first considered only validation data from our study area (Model 2), which produced slightly lower point estimates for HF183 sensitivity (60%) than obtained from crude calculations (64%). The sensitivity estimates were similar for all three sample types, with the greatest uncertainty in stored water (95% CI: 38-81%). The HF183 specificity estimate was similar to the crude value (67%) for latrine soil [66% (53-80%)] but the estimates were higher for household soil [72% (61-83%)] and stored water [85% (77-91%)], in which HF183 was detected less frequently. Sensitivity and specificity patterns for Mnif followed similar patterns.

Meta-analytic sensitivity and specificity estimates were slightly higher than the single-study estimates when using a single indicator (Model 3) and when combining both indicators (Model 4). However, the 95% CIs remained similar for all three models, suggesting the inclusion of additional information minimally impacted estimates of diagnostic accuracy. Model 5, which simultaneously considered both indicators in all sample types, provided a single set of sensitivity and specificity estimates for each indictor. The estimates were comparable to values observed for individual sample types but featured narrower 95% CIs, indicating reduced uncertainty in indicator performance with the inclusion of data from multiple sample types. While the estimated sensitivity for Mnif [71% (59-83%)] remained similar to the crude value (71%), the estimated specificity of both indicators and HF183 sensitivity were somewhat higher than expected from crude calculations.

**S11. Accuracy-adjusted effects**

We constructed a series of models to estimate sanitation intervention effects on the prevalence of human fecal contamination. In the first model, human-associated fecal indicators were used as direct proxies for human fecal contamination, equivalent to assuming 100% sensitivity and specificity. The estimated intervention effect, given by the DID prevalence odds ratio, was less than one for both indicators in latrine soil and household soil and above one for HF183 in stored water (Figure S1). However, the estimates were imprecise and encompassed both large increases and large reductions within the 95% CIs, providing little evidence for an effect of the intervention on either indicator in any sample type. Incorporating local validation data to account for indicator accuracy (Model 2) further widened the 95% CIs, indicating additional uncertainty about the intervention effect on the prevalence of human fecal contamination. Effect estimates incorporating multiple validation datasets (Model 3) were largely similar to those using local validation data alone. When using HF183 and Mnif to jointly estimate human fecal contamination (Model 4), the DID estimates were similar to estimates for individual indicators. Similarly, combining observations from all three sample types produced an estimated intervention effect close to the null with the 95% CI encompassing both reductions and increases in the odds of human fecal contamination at the compound level [POR: 0.93 (95% CI: 0.42-2.1)]. Across all model formulations, there was little evidence for an effect of the sanitation intervention on the prevalence of human fecal contamination.

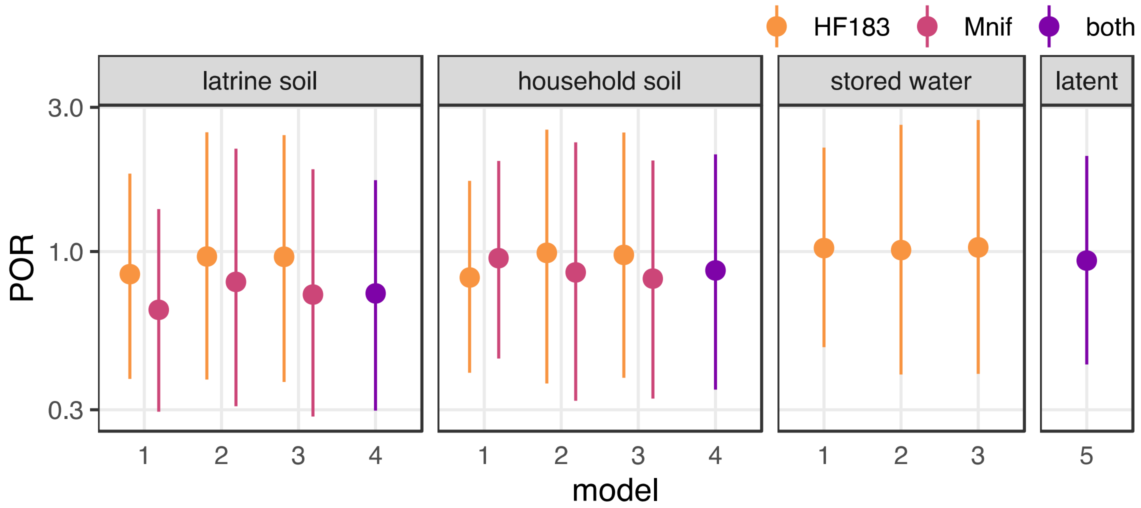

**Figure S1. Difference-in-difference prevalence odds ratio (POR) estimates of the sanitation intervention effect on human fecal contamination under five different models. Model 1 made no correction for indicator accuracy, Model 2 used local validation data to account for indicator sensitivity and specificity, and Models 3, 4 and 5 used meta-analytic estimates of local indicator accuracy. Model 3 was fit separately by indicator and sample type, Model 4 was fit to both indicators by sample type, and Model 5 used both indicators in all sample types to estimate the latent compound prevalence of human fecal contamination. All models were adjusted for population density, presence of animals, wealth score, temperature, antecedent precipitation, and sun exposure and surface wetness for soil samples and storage container mouth width and cover status for water samples.**

### S12. Human fecal contamination prevalence estimates

Posterior predictions from each of the five models were used to estimate stratum-specific prevalence. We fit unadjusted models that included only intercepts and the DID terms (Table S10) as well as adjusted models with meteorological, compound, household, and sample characteristics included as covariates (Table 2, main text).

**Table S10. Estimated sensitivity, specificity, and prevalence of human fecal contamination from unadjusted models**

| | | | | | prevalence estimate (95% CI) | | | | |
|---|---|---|---|---|---|---|---|---|---|
| | | | | | control | | intervention | | |
| | target | sensitivity (95% CI) | specificity (95% CI) | N | before | after | before | after | prevalence DID (95% CI) |
| **latrine soil** | | | | | | | | | |
| bootstrap | HF183 | 1 | 1 | 116 | 0.33 (0.17, 0.50) | 0.57 (0.39, 0.75) | 0.43 (0.23, 0.64) | 0.43 (0.26, 0.61) | -0.23 (-0.60, 0.14) |
| | Mnif | 1 | 1 | 116 | 0.51 (0.35, 0.69) | 0.50 (0.32, 0.68) | 0.65 (0.45, 0.84) | 0.36 (0.19, 0.54) | -0.27 (-0.63, 0.08) |
| model 1 | HF183 | 1 | 1 | 116 | 0.41 (0.28, 0.54) | 0.49 (0.35, 0.64) | 0.40 (0.26, 0.56) | 0.45 (0.30, 0.61) | -0.04 (-0.22, 0.14) |
| | Mnif | 1 | 1 | 116 | 0.54 (0.41, 0.67) | 0.48 (0.35, 0.62) | 0.57 (0.41, 0.71) | 0.42 (0.27, 0.57) | -0.09 (-0.27, 0.09) |
| model 2 | HF183 | 0.59 (0.41, 0.80) | 0.65 (0.52, 0.79) | 116 | 0.41 (0.05, 0.90) | 0.44 (0.05, 0.92) | 0.41 (0.04, 0.91) | 0.43 (0.04, 0.93) | -0.01 (-0.20, 0.19) |
| | Mnif | 0.68 (0.50, 0.86) | 0.67 (0.52, 0.82) | 116 | 0.53 (0.11, 0.92) | 0.48 (0.09, 0.91) | 0.53 (0.10, 0.92) | 0.44 (0.07, 0.93) | -0.05 (-0.25, 0.14) |
| model 3 | HF183 | 0.65 (0.45, 0.85) | 0.68 (0.55, 0.83) | 116 | 0.39 (0.06, 0.87) | 0.43 (0.06, 0.89) | 0.38 (0.05, 0.88) | 0.41 (0.04, 0.91) | -0.01 (-0.19, 0.18) |
| | Mnif | 0.70 (0.55, 0.83) | 0.70 (0.55, 0.84) | 116 | 0.54 (0.17, 0.87) | 0.49 (0.13, 0.88) | 0.55 (0.14, 0.88) | 0.44 (0.09, 0.88) | -0.05 (-0.25, 0.14) |
| model 4 | HF183 | 0.64 (0.45, 0.84) | 0.69 (0.55, 0.83) | 116 | 0.47 (0.16, 0.79) | 0.46 (0.13, 0.79) | 0.47 (0.13, 0.81) | 0.40 (0.10, 0.77) | -0.06 (-0.25, 0.13) |
| | Mnif | 0.71 (0.56, 0.84) | 0.69 (0.54, 0.83) | | | | | | |
| model 5 | HF183 | 0.72 (0.55, 0.87) | 0.83 (0.76, 0.90) | 107 | 0.48 (0.25, 0.73) | 0.49 (0.27, 0.72) | 0.43 (0.19, 0.70) | 0.47 (0.25, 0.71) | 0.04 (-0.14, 0.23) |
| | Mnif | 0.71 (0.58, 0.84) | 0.79 (0.70, 0.88) | | | | | | |
| **household soil** | | | | | | | | | |
| bootstrap | HF183 | 1 | 1 | 176 | 0.17 (0.07, 0.28) | 0.49 (0.35, 0.64) | 0.36 (0.20, 0.52) | 0.38 (0.24, 0.52) | -0.30 (-0.57, -0.01) |
| | Mnif | 1 | 1 | 175 | 0.43 (0.30, 0.57) | 0.25 (0.13, 0.39) | 0.23 (0.09, 0.37) | 0.24 (0.12, 0.38) | 0.20 (-0.07, 0.46) |
| model 1 | HF183 | 1 | 1 | 176 | 0.27 (0.18, 0.37) | 0.41 (0.30, 0.53) | 0.31 (0.20, 0.44) | 0.40 (0.28, 0.53) | -0.05 (-0.21, 0.11) |
| | Mnif | 1 | 1 | 175 | 0.37 (0.26, 0.48) | 0.29 (0.19, 0.40) | 0.29 (0.18, 0.41) | 0.24 (0.14, 0.35) | 0.03 (-0.11, 0.17) |
| model 2 | HF183 | 0.58 (0.36, 0.80) | 0.71 (0.61, 0.83) | 176 | 0.25 (0.03, 0.78) | 0.30 (0.02, 0.81) | 0.26 (0.02, 0.78) | 0.31 (0.02, 0.84) | 0.00 (-0.18, 0.19) |
| | Mnif | 0.65 (0.41, 0.86) | 0.75 (0.66, 0.85) | 175 | 0.22 (0.03, 0.55) | 0.18 (0.02, 0.48) | 0.18 (0.02, 0.49) | 0.13 (0.01, 0.43) | 0.00 (-0.13, 0.13) |
| model 3 | HF183 | 0.65 (0.40, 0.85) | 0.74 (0.63, 0.85) | 176 | 0.23 (0.03, 0.64) | 0.30 (0.03, 0.71) | 0.23 (0.03, 0.66) | 0.30 (0.02, 0.76) | 0.00 (-0.18, 0.18) |
| | Mnif | 0.69 (0.52, 0.84) | 0.77 (0.68, 0.86) | 175 | 0.22 (0.03, 0.50) | 0.17 (0.02, 0.41) | 0.17 (0.02, 0.41) | 0.12 (0.01, 0.34) | 0.00 (-0.12, 0.13) |
| model 4 | HF183 | 0.67 (0.44, 0.86) | 0.71 (0.62, 0.80) | 175 | 0.15 (0.02, 0.35) | 0.16 (0.02, 0.39) | 0.13 (0.02, 0.31) | 0.13 (0.01, 0.34) | -0.01 (-0.12, 0.11) |
| | Mnif | 0.69 (0.52, 0.83) | 0.76 (0.67, 0.85) | | | | | | |
| model 5 | HF183 | 0.72 (0.55, 0.87) | 0.83 (0.76, 0.90) | 156 | 0.24 (0.09, 0.43) | 0.24 (0.10, 0.42) | 0.20 (0.06, 0.39) | 0.23 (0.08, 0.42) | 0.03 (-0.11, 0.17) |
| | Mnif | 0.71 (0.58, 0.84) | 0.79 (0.70, 0.88) | | | | | | |
| **stored water** | | | | | | | | | |
| bootstrap | HF183 | 1 | 1 | 193 | 0.12 (0.04, 0.22) | 0.10 (0.02, 0.20) | 0.22 (0.10, 0.35) | 0.19 (0.09, 0.30) | -0.01 (-0.21, 0.19) |
| model 1 | HF183 | 1 | 1 | 193 | 0.14 (0.08, 0.23) | 0.12 (0.06, 0.20) | 0.19 (0.11, 0.29) | 0.18 (0.10, 0.28) | 0.01 (-0.09, 0.11) |
| model 2 | HF183 | 0.58 (0.34, 0.80) | 0.84 (0.78, 0.89) | 193 | 0.08 (0.01, 0.23) | 0.07 (0.01, 0.22) | 0.08 (0.01, 0.27) | 0.07 (0.01, 0.26) | 0.00 (-0.08, 0.09) |
| model 3 | HF183 | 0.65 (0.40, 0.85) | 0.85 (0.79, 0.90) | 193 | 0.07 (0.01, 0.20) | 0.06 (0.01, 0.19) | 0.08 (0.01, 0.25) | 0.07 (0.01, 0.24) | 0.00 (-0.08, 0.08) |
| model 5 | HF183 | 0.72 (0.55, 0.87) | 0.83 (0.76, 0.90) | 176 | 0.08 (0.01, 0.20) | 0.09 (0.01, 0.21) | 0.07 (0.01, 0.18) | 0.08 (0.01, 0.20) | 0.01 (-0.05, 0.08) |
| **latent compound** | | | | | | | | | |
| m5 | HF183 | 0.72 (0.55, 0.87) | 0.83 (0.76, 0.90) | 113 | 0.26 (0.08, 0.55) | 0.26 (0.08, 0.55) | 0.22 (0.06, 0.50) | 0.25 (0.07, 0.53) | 0.03 (-0.11, 0.18) |
| | Mnif | 0.71 (0.58, 0.84) | 0.79 (0.70, 0.88) | | | | | | |

### S13. Prior distributions

For all models, we sought to select regularizing ("weakly informative") priors where feasible. Regularizing priors impose soft constraints on parameter values, discouraging the model fitting algorithm from exploring extreme values that can reasonably be assumed implausible *a priori*.[25] In addition to the practical benefit of often aiding model convergence, particularly for complex, high-dimensional models, regularization can help improve the precision of estimates for parameters for which the underlying data are somewhat noisy. This also has the effect of shrinking estimates towards a common value—often the null, in the case of regression coefficients, which mildly increases the strength of the evidence necessary to demonstrate a probable effect but provides the advantage of reducing false positives that can arise from typical sampling variation.[25] While setting regularizing priors is subjective in the strictest sense, one generally possesses sufficient information to determine a broadly plausible range of parameter values that arguably are more readily accepted than the assumption implicit in flat ("non-informative") priors that the range of potential parameter values is essentially infinite.

For the basic DID models used to assess the intervention effect on individual fecal indicators in separate sample types (Table S8), we included a compound-varying intercept that used the **brms** package default positive-constrained student-t prior with 3 degrees of freedom (df) and scale determined from the link-transformed data.[39] Censored linear regression was used to estimate the intervention impact on the $\log_{10}$ concentrations of the two *E. coli* assays, cEC and EC23S, using regularizing normal priors with standard deviation (SD) = 5 on the population-level intercept and SD = 2 on the predictor coefficients, including the DID terms.[25,39,40] We estimated the effect of the intervention on human-associated indicator prevalence using logistic regression and the prevalence odds ratio (POR) as the measure of effect. Under the logistic link, priors are defined on the continuous log-odds scale but are best understood on the probability scale, which is more intuitive but constrained between 0 and 1 prevalence.[41] As such, the population-level intercept and predictor coefficients were given regularizing normal priors with SD = 1.5 and SD = 0.5, respectively, which on the probability scale corresponded to a ~95% chance the population-level prevalence was between 0.05 and 0.95 and an effect of up to $\pm 0.23$ for each predictor at a population-level prevalence of 0.5.[40] This represents a substantial effect size on the probability scale—a reduction in absolute risk from 50% to 27%, for example—which would be unexpected in an environmental context.[23] Furthermore, as a soft constraint, estimates could still exceed this effect size given sufficient and sufficiently strong data in support of a larger effect.

The diagnostic accuracy-corrected models of human fecal contamination prevalence shared the same basic structure as the human-associated indicator prevalence model described above. Accordingly, we set the same priors for the population-level intercept, the DID terms, and the covariates in the adjustment set:

$$y_i \sim Bernoulli(p_i)$$

$$p_i = \pi_i$$

$$logit(\pi_i) = \beta^0 + \beta^P P_i + \beta^T T_i + \beta^{DID} P_i \times T_i + \boldsymbol{X}_i \boldsymbol{\gamma}$$ **Model 1**

$$\beta^0 \sim normal(0,1.5)$$

$$\beta^P, \beta^T, \beta^{DID}, \boldsymbol{\gamma} \sim normal(0,0.5)$$

Model 2 introduced parameters for sensitivity ($Se$) and specificity ($Sp$), which required priors as well. Due the modest sample sizes of the local validation analysis, particularly of human samples (n=14), the use of non-informative priors risks unrealistically broad sensitivity and specificity estimates.[23,42] As we possessed prior information on the typical ranges for $Se$ and

$Sp$ in resource-limited settings (Table S9), we used informative $beta(3,2)$ priors as soft constraints, corresponding to a 95% chance they fall between 0.19 and 0.93 on the probability scale, with mean 0.6.

$$y_i \sim Bernoulli(p_i)$$
$$p_i = Se \times \pi_i + (1 - Sp)(1 - \pi_i)$$
$$logit(\pi_i) = \beta^0 + \beta^P P_i + \beta^T T_i + \beta^{DID} P_i \times T_i + \boldsymbol{X}_i \boldsymbol{\gamma}$$
$$y^{Se} \sim binomial(n^{Se}, Se) \qquad \textbf{Model 2}$$
$$y^{Sp} \sim binomial(n^{Sp}, Sp)$$
$$Se, Sp \sim beta(3,2)$$
$$\beta^0 \sim normal(0,1.5);\ \beta^P, \beta^T, \beta^{DID}, \boldsymbol{\gamma} \sim normal(0,0.5)$$

Because our validation sample set was small and performance estimates vary widely between studies, we fit a third model (Model 3) featuring a meta-analysis of indicator sensitivity and specificity. We assumed the log-odds of the sensitivity in the $k$th study, $Se_{[k]}$, were normally distributed with mean $\mu^{Se}$ and SD $\sigma^{Se}$, with an equivalent structure for specificity. We assigned $\mu^{Se}$ and $\mu^{Sp}$ $normal(0.5, 1)$ priors, which provided approximately equivalent coverage on the probability scale as the previous beta priors in Model 2, with weakly-informative $normal^{+}(0, 0.5)$ [half-normal] priors on $\sigma^{Se}$ and $\sigma^{Sp}$, as discussed by Gelman and Carpenter.[23]

$$y_i \sim Bernoulli(p_i)$$

$$p_i = Se_{[1]} \times \pi_i + (1 - Sp_{[1]})(1 - \pi_i)$$

$$logit(\pi_i) = \beta^0 + \beta^P P_i + \beta^T T_i + \beta^{DID} P_i \times T_i + \boldsymbol{X}_i \boldsymbol{\gamma}$$

$$y_{[k]}^{Se} \sim binomial(n_{[k]}^{Se}, Se_{[k]});\ y_{[k]}^{Sp} \sim binomial(n_{[k]}^{Sp}, Sp_{[k]})$$

**Model 3**

$$logit(Se_{[k]}) \sim normal(\mu^{Se}, \sigma^{Se});\ logit(Sp_{[k]}) \sim normal(\mu^{Sp}, \sigma^{Sp})$$

$$\mu^{Se}, \mu^{Sp} \sim normal(0.5, 1)$$

$$\sigma^{Se}, \sigma^{Sp} \sim normal^{+}(0, 0.5)$$

$$\beta^0 \sim normal(0,1.5);\ \beta^P, \beta^T, \beta^{DID}, \boldsymbol{\gamma} \sim normal(0,0.5)$$

Model 4 followed the same structure as Model 3, but incorporated two different fecal indicators, represented as $y_i^{[assay]}$, with a corresponding duplication of the sensitivity and specificity model components. All priors remain the same as Model 3.

$$y_i^{[assay]} \sim Bernoulli\left(p_i^{[assay]}\right)$$

$$p_i^{[assay]} = Se_{[1]}^{[assay]} \times \pi_i + \left(1 - Sp_{[1]}^{[assay]}\right)(1 - \pi_i)$$

$$logit(\pi_i) = \beta^0 + \beta^P P_i + \beta^T T_i + \beta^{DID} P_i \times T_i + \boldsymbol{X}_i \boldsymbol{\gamma}$$

$$y_{[k]}^{[assay],Se} \sim binomial\left(n_{[k]}^{[assay],Se,}, Se_{[k]}^{[assay]}\right);\ y_{[k]}^{[assay],Sp} \sim binomial\left(n_{[k]}^{[assay],Sp,}, Sp_{[k]}^{[assay]}\right)$$

$$logit\left(Se_{[k]}^{[assay]}\right) \sim normal\left(\mu^{[assay],Se}, \sigma^{[assay],Se}\right)$$

**Model 4**

$$logit\left(Sp_{[k]}^{[assay]}\right) \sim normal\left(\mu^{[assay],Sp}, \sigma^{[assay],Sp}\right)$$

$$\mu^{[assay],Se}, \mu^{[assay],Sp} \sim normal(0.5, 1)$$

$$\sigma^{[assay],Se}, \sigma^{[assay],Sp} \sim normal^{+}(0, 0.5)$$

$$\beta^0 \sim normal(0,1.5);\ \beta^P, \beta^T, \beta^{DID}, \boldsymbol{\gamma} \sim normal(0,0.5)$$

Finally, Model 5 included multiple sample types with type-specific prevalence variables, $\pi_i^{[type]}$, derived from sample-type specific intercepts $\alpha^{[type]}$ and a shared compound-varying intercept $\alpha_{[j]}^{comp}$ corresponding to the $j$the compound that replaced the previously fixed, population-level intercept $\beta^0$. As before, all $\beta$ and $\gamma$ parameters were given normal priors with SD = 0.5. We assumed $\alpha_{[j]}^{comp} \sim normal(\mu^{comp}, \sigma^{comp})$ and $\alpha^{[type]} \sim normal(0, \sigma^{type})$, with half-normal, SD = 0.5 priors on $\sigma^{comp}$ and $\sigma^{type}$ and the same $normal(0, 1.5)$ prior on $\mu^{comp}$ used for the fixed intercept $\beta^0$ in previous models.

$$y_i^{[assay,type]} \sim Bernoulli\left(p_i^{[assay,type]}\right)$$

$$p_i^{[assay,type]} = Se_{[1]}^{[assay,type]} \times \pi_i^{[type]} + \left(1 - Sp_{[1]}^{[assay,type]}\right)\left(1 - \pi_i^{[type]}\right)$$

$$logit(\pi_i^{[type]}) = \alpha^{[type]} + \boldsymbol{X}_i^{[type]}\boldsymbol{\gamma}^{[type]} + logit\left(\pi_{[j]}^{comp}\right)$$

$$logit\left(\pi_{[j]}^{comp}\right) = \alpha_{[j]}^{comp} + \beta^P P_{[j]} + \beta^T T_{[j]} + \beta^{DID} P_{[j]} \times T_{[j]} + \boldsymbol{X}_{[j]}^{comp}\boldsymbol{\gamma}^{comp}$$

$$y_{[k]}^{[assay],Se} \sim binomial\left(n_{[k]}^{[assay],Se,}, Se_{[k]}^{[assay]}\right);\ y_{[k]}^{[assay],Sp} \sim binomial\left(n_{[k]}^{[assay],Sp,}, Sp_{[k]}^{[assay]}\right)$$

**Model 5**

$$logit\left(Se_{[k]}^{[assay]}\right) \sim normal\left(\mu^{[assay],Se}, \sigma^{[assay],Se}\right)$$

$$logit\left(Sp_{[k]}^{[assay]}\right) \sim normal\left(\mu^{[assay],Sp}, \sigma^{[assay],Sp}\right)$$

$$\alpha_{[j]}^{comp} \sim normal(\mu^{comp}, \sigma^{comp});\ \alpha^{[type]} \sim normal(0, \sigma^{type})$$

$$\mu^{[assay],Se}, \mu^{[assay],Sp} \sim normal(0.5, 1);\ \mu^{comp} \sim normal(0,1.5)$$

$$\sigma^{[type]}, \sigma^{[assay],Se}, \sigma^{[assay],Sp} \sim normal^+(0, 0.5)$$

$$\beta^P, \beta^T, \beta^{DID}, \boldsymbol{\gamma}^{[type]}, \boldsymbol{\gamma}^{comp} \sim normal(0,0.5)$$

### S14. Stan code

#### *S14.1. Model 1*

```
// Model 1:
// single sample type, single target
// no correction for sensitivity/specificity
// no compound-varying intercepts

data{
// sample data
  int<lower = 0> N_samp;  // number of sample observations
  int<lower = 0, upper = 1> y_samp[N_samp];  // sample observations
  vector<lower = 0, upper = 1>[N_samp] phase;  // survey phase
  vector<lower = 0, upper = 1>[N_samp] treat;  // treatment arm
  vector<lower = 0, upper = 1>[N_samp] did;  // phase*treat interaction
  int<lower = 0> K;  // number of predictors
  matrix[N_samp, K] X;  // predictor values

// prior predictive check?
  int<lower = 0, upper = 1> prior_only;  // toggle on to sample priors only
}

parameters{
// linear model parameters
  real b0;  // intercept
  real bP;  // phase
  real bT;  // treatment
  real bD;  // DID
  vector[K] g;  // predictor coefficients
}

model{
// linear model for probability of human contamination
  vector[N_samp] p_samp = inv_logit(b0 + bP*phase + bT*treat + bD*did + X*g);

// Likelihood
  if(prior_only == 0){
    y_samp ~ binomial(1, p_samp);
  }

// linear model priors
  b0 ~ normal(0, 1.5);
  bD ~ normal(0, 0.5);
  bT ~ normal(0, 0.5);
  bP ~ normal(0, 0.5);
  g ~ normal(0, 0.5);
}

generated quantities{
// posterior predictions (or prior predictions, if prior_only == 1)
// define predicted variables
  int<lower = 0, upper = 1> y_pred[N_samp];  // predicted sample observations
  int<lower = 0> n_pos;  // number of predicted positives
  real<lower = 0, upper = 1> p_samp_avg;  // mean target prevalence in samples
```

```
// calculate human contamination probability
  vector[N_samp] p_samp_sim = inv_logit(b0 + bP*phase + bT*treat + bD*did + X*g);

// predict sample observations
  y_pred = binomial_rng(1, p_samp_sim);
  n_pos = sum(y_pred);

// summarise prevalence calculations
  p_samp_avg = mean(p_samp_sim);
}
```

*S14.2. Model 2*

```
// Model 2:
// single sample type, single target
// diagnostic performance data from this study only
// no compound-varying intercepts

data{
// sample data
  int<lower = 0> N_samp;  // number of sample observations
  int<lower = 0, upper = 1> y_samp[N_samp];  // sample observations
  vector<lower = 0, upper = 1>[N_samp] phase;  // survey phase
  vector<lower = 0, upper = 1>[N_samp] treat;  // treatment arm
  vector<lower = 0, upper = 1>[N_samp] did;  // phase*treat interaction
  int<lower = 0> K;  // number of predictors
  matrix[N_samp, K] X;  // predictor values

// sens/spec data
  int<lower = 0> y_spec;  // number of true negatives observed
  int<lower = 0> n_spec;  // number of non-target samples
  int<lower = 0> y_sens;  // number of true positives observed
  int<lower = 0> n_sens;  // number of target samples

// prior predictive check?
  int<lower = 0, upper = 1> prior_only;  // toggle on to sample priors only
}

parameters{
// linear model parameters
  real b0;  // intercept
  real bP;  // phase
  real bT;  // treatment
  real bD;  // DID
  vector[K] g;  // predictor coefficients

// diagnostic performance parameters
  real<lower=0, upper=1> spec;  // specificity
  real<lower=0, upper=1> sens;  // sensitivity
}

model{
// linear model for probability of human contamination
  vector[N_samp] p = inv_logit(b0 + bP*phase + bT*treat + bD*did + X*g);

// adjust for sens/spec
  vector[N_samp] p_samp = sens * p + (1 - spec) * (1 - p);

// Likelihoods
  if(prior_only == 0){
  // samples
    y_samp ~ binomial(1, p_samp);

  // validation studies
    y_spec ~ binomial(n_spec, spec);
    y_sens ~ binomial(n_sens, sens);
  }
```

```
// linear model priors
  b0 ~ normal(0, 1.5);
  bD ~ normal(0, 0.5);
  bT ~ normal(0, 0.5);
  bP ~ normal(0, 0.5);
  g ~ normal(0, 0.5);

// validation priors
  sens ~ beta(3, 2);
  spec ~ beta(3, 2);
}

generated quantities{
// posterior predictions (or prior predictions, if prior_only == 1)
// define predicted variables
  int<lower = 0, upper = 1> y_pred[N_samp];  // predicted sample observations
  int<lower = 0> n_pos;  // number of predicted positives
  real<lower = 0, upper = 1> p_avg;  //  mean human contamination prevalence
  real<lower = 0, upper = 1> p_samp_avg;  // mean target prevalence in samples

// calculate human contamination probability
  vector[N_samp] p_sim = inv_logit(b0 + bP*phase + bT*treat + bD*did + X*g);

// adjust for sens/spec
  vector[N_samp] p_samp_sim = sens * p_sim + (1 - spec) * (1 - p_sim);

// predict sample observations
  y_pred = binomial_rng(1, p_samp_sim);
  n_pos = sum(y_pred);

// summarise prevalence calculations
  p_avg = mean(p_sim);
  p_samp_avg = mean(p_samp_sim);
}
```

*S14.3. Model 3*

```
// Model 3:
// single sample type, single target
// diagnostic performance meta-analysis
// no compound-varying intercepts

data{
// sample data
  int<lower = 0> N_samp;  // number of sample observations
  int<lower = 0, upper = 1> y_samp[N_samp];  // sample observations
  vector<lower = 0, upper = 1>[N_samp] phase;  // survey phase
  vector<lower = 0, upper = 1>[N_samp] treat;  // treatment arm
  vector<lower = 0, upper = 1>[N_samp] did;  // phase*treat interaction
  int<lower = 0> K;  // number of predictors
  matrix[N_samp, K] X;  // predictor values

// sens/spec data
  int<lower = 0> J_spec;
  int<lower = 0> y_spec[J_spec];
  int<lower = 0> n_spec[J_spec];
  int<lower = 0> J_sens;
  int<lower = 0> y_sens[J_sens];
  int<lower = 0> n_sens[J_sens];

// prior predictive check?
  int<lower = 0, upper = 1> prior_only;  // toggle on to sample priors only
}

parameters{
// linear model parameters
  real b0;  // intercept
  real bP;  // phase
  real bT;  // treatment
  real bD;  // DID
  vector[K] g;  // predictor coefficients

// sens/spec meta-analysis parameters
  real mu_logit_spec;  // mean spec on the logit scale
  real mu_logit_sens;
  real<lower = 0> sigma_logit_spec;  // spec SD on logit scale
  real<lower = 0> sigma_logit_sens;
  // non-centered parameterization of logit-transformed sens/spec
  vector<offset = mu_logit_spec, multiplier = sigma_logit_spec>[J_spec] logit_spec;
  vector<offset = mu_logit_sens, multiplier = sigma_logit_sens>[J_sens] logit_sens;
}

transformed parameters{
// recover sens/spec on probability scale
  vector[J_spec] spec = inv_logit(logit_spec);
  vector[J_sens] sens = inv_logit(logit_sens);
}

model{
// linear model for probability of human contamination
  vector[N_samp] p = inv_logit(b0 + bP*phase + bT*treat + bD*did + X*g);
```

```
// adjust for sens/spec
  vector[N_samp] p_samp = sens[1] * p + (1 - spec[1]) * (1 - p);

// Likelihoods
  if(prior_only == 0){
  // samples
    y_samp ~ binomial(1, p_samp);

  // validation studies
    y_spec ~ binomial(n_spec, spec);
    y_sens ~ binomial(n_sens, sens);
  }

// linear model priors
  b0 ~ normal(0, 1.5);
  bD ~ normal(0, 0.5);
  bT ~ normal(0, 0.5);
  bP ~ normal(0, 0.5);
  g ~ normal(0, 0.5);

// validation priors
  logit_spec ~ normal(mu_logit_spec, sigma_logit_spec);
  logit_sens ~ normal(mu_logit_sens, sigma_logit_sens);
  sigma_logit_spec ~ normal(0, .5);
  sigma_logit_sens ~ normal(0, .5);
  mu_logit_spec ~ normal(.5, 1);
  mu_logit_sens ~ normal(.5, 1);
}

generated quantities{
// posterior predictions (or prior predictions, if prior_only == 1)
// define predicted variables
  int<lower = 0, upper = 1> y_pred[N_samp];  // predicted sample observations
  int<lower = 0> n_pos;  // number of predicted positives
  real<lower = 0, upper = 1> p_avg;  //  mean human contamination prevalence
  real<lower = 0, upper = 1> p_samp_avg;  // mean target prevalence in samples

// calculate human contamination probability
  vector[N_samp] p_sim = inv_logit(b0 + bP*phase + bT*treat + bD*did + X*g);

// adjust for sens/spec
  vector[N_samp] p_samp_sim = sens[1] * p_sim + (1 - spec[1]) * (1 - p_sim);

// predict sample observations
  y_pred = binomial_rng(1, p_samp_sim);
  n_pos = sum(y_pred);

// summarise prevalence calculations
  p_avg = mean(p_sim);
  p_samp_avg = mean(p_samp_sim);
}
```

*S14.4. Model 4*

```
// Model 4:
// single sample type, two targets
// diagnostic performance meta-analysis
// no compound-varying intercepts

data{
// sample data
  int<lower = 0> N_samp;  // number of sample observations
  int<lower = 0, upper = 1> y_hf[N_samp];  // HF183 observations
  int<lower = 0, upper = 1> y_mn[N_samp];  // Mnif observations
  vector<lower = 0, upper = 1>[N_samp] phase;  // survey phase
  vector<lower = 0, upper = 1>[N_samp] treat;  // treatment arm
  vector<lower = 0, upper = 1>[N_samp] did;  // phase*treat interaction
  int<lower = 0> K;  // number of predictors
  matrix[N_samp, K] X;  // predictor values

// sens/spec data
  int<lower = 0> J_spec_hf;
  int<lower = 0> y_spec_hf[J_spec_hf];
  int<lower = 0> n_spec_hf[J_spec_hf];
  int<lower = 0> J_sens_hf;
  int<lower = 0> y_sens_hf[J_sens_hf];
  int<lower = 0> n_sens_hf[J_sens_hf];
  int<lower = 0> J_spec_mn;
  int<lower = 0> y_spec_mn[J_spec_mn];
  int<lower = 0> n_spec_mn[J_spec_mn];
  int<lower = 0> J_sens_mn;
  int<lower = 0> y_sens_mn[J_sens_mn];
  int<lower = 0> n_sens_mn[J_sens_mn];

// prior predictive check?
  int<lower = 0, upper = 1> prior_only;  // toggle on to sample priors only
}

parameters{
// linear model parameters
  real b0;  // intercept
  real bP;  // phase
  real bT;  // treatment
  real bD;  // DID
  vector[K] g;  // predictor coefficients

// sens/spec meta-analysis parameters
  real mu_logit_spec_hf;  // mean spec for HF183 on the logit scale
  real mu_logit_sens_hf;
  real mu_logit_spec_mn;
  real mu_logit_sens_mn;
  real<lower = 0> sigma_logit_spec_hf;  // spec SD on logit scale
  real<lower = 0> sigma_logit_sens_hf;
  real<lower = 0> sigma_logit_spec_mn;
  real<lower = 0> sigma_logit_sens_mn;
  // non-centered parameterization of logit-transformed sens/spec for each target
  vector<offset = mu_logit_spec_hf, multiplier = sigma_logit_spec_hf>[J_spec_hf]
logit_spec_hf;
```

```
  vector<offset = mu_logit_sens_hf, multiplier = sigma_logit_sens_hf>[J_sens_hf]
logit_sens_hf;
  vector<offset = mu_logit_spec_mn, multiplier = sigma_logit_spec_mn>[J_spec_mn]
logit_spec_mn;
  vector<offset = mu_logit_sens_mn, multiplier = sigma_logit_sens_mn>[J_sens_mn]
logit_sens_mn;
}

transformed parameters{
// recover sens/spec on probability scale
  vector[J_spec_hf] spec_hf = inv_logit(logit_spec_hf);
  vector[J_sens_hf] sens_hf = inv_logit(logit_sens_hf);
  vector[J_spec_mn] spec_mn = inv_logit(logit_spec_mn);
  vector[J_sens_mn] sens_mn = inv_logit(logit_sens_mn);
}

model{
// linear model for probability of human contamination
  vector[N_samp] p = inv_logit(b0 + bP*phase + bT*treat + bD*did + X*g);

// adjust for sens/spec
  // by convention the first sens/spec element represents this current study
  vector[N_samp] p_hf = sens_hf[1] * p + (1 - spec_hf[1]) * (1 - p);
  vector[N_samp] p_mn = sens_mn[1] * p + (1 - spec_mn[1]) * (1 - p);

// Likelihoods
  if(prior_only == 0){
  // samples
    y_hf ~ binomial(1, p_hf);
    y_mn ~ binomial(1, p_mn);

  // validation studies
    y_spec_hf ~ binomial(n_spec_hf, spec_hf);
    y_sens_hf ~ binomial(n_sens_hf, sens_hf);
    y_spec_mn ~ binomial(n_spec_mn, spec_mn);
    y_sens_mn ~ binomial(n_sens_mn, sens_mn);
  }

// linear model priors
  b0 ~ normal(0, 1.5);
  bD ~ normal(0, 0.5);
  bT ~ normal(0, 0.5);
  bP ~ normal(0, 0.5);
  g ~ normal(0, 0.5);

// validation priors
  logit_spec_hf ~ normal(mu_logit_spec_hf, sigma_logit_spec_hf);
  logit_sens_hf ~ normal(mu_logit_sens_hf, sigma_logit_sens_hf);
  sigma_logit_spec_hf ~ normal(0, .5);
  sigma_logit_sens_hf ~ normal(0, .5);
  mu_logit_spec_hf ~ normal(.5, 1);
  mu_logit_sens_hf ~ normal(.5, 1);
  logit_spec_mn ~ normal(mu_logit_spec_mn, sigma_logit_spec_mn);
  logit_sens_mn ~ normal(mu_logit_sens_mn, sigma_logit_sens_mn);
  sigma_logit_spec_mn ~ normal(0, .5);
  sigma_logit_sens_mn ~ normal(0, .5);
  mu_logit_spec_mn ~ normal(.5, 1);
```

```
  mu_logit_sens_mn ~ normal(.5, 1);
}

generated quantities{
// posterior predictions (or prior predictions, if prior_only == 1)
// define predicted variables
  int<lower = 0, upper = 1> y_pred_hf[N_samp];  // predicted HF183 observations
  int<lower = 0> n_pos_hf;  // number of predicted HF183 positives
  real<lower = 0, upper = 1> p_samp_avg_hf;  // mean HF183 prevalence in samples
  int<lower = 0, upper = 1> y_pred_mn[N_samp];  // predicted Mnif observations
  int<lower = 0> n_pos_mn;  // number of predicted Mnif positives
  real<lower = 0, upper = 1> p_samp_avg_mn;  // mean Mnif prevalence in samples
  real<lower = 0, upper = 1> p_avg;  //  mean human contamination prevalence

// calculate human contamination probability
  vector[N_samp] p_sim = inv_logit(b0 + bP*phase + bT*treat + bD*did + X*g);

// adjust for sens/spec
  vector[N_samp] p_hf_sim = sens_hf[1] * p_sim + (1 - spec_hf[1]) * (1 - p_sim);
  vector[N_samp] p_mn_sim = sens_mn[1] * p_sim + (1 - spec_mn[1]) * (1 - p_sim);

// predict sample observations
  y_pred_hf = binomial_rng(1, p_hf_sim);
  n_pos_hf = sum(y_pred_hf);
  y_pred_mn = binomial_rng(1, p_mn_sim);
  n_pos_mn = sum(y_pred_mn);

// summarise prevalence calculations
  p_avg = mean(p_sim);
  p_samp_avg_hf = mean(p_hf_sim);
  p_samp_avg_mn = mean(p_mn_sim);
}
```

*S14.5. Model 5*

```
// Model 5:
// three sample types, two targets
// diagnostic performance meta-analysis
// compound-varying intercept, type-varying intercept

data{
  int<lower = 0> N_type;  // number of sample types considered

// compound data
  int<lower = 0> J_comp;  // number of unique compounds
  int<lower = 0> N_comp;  // number of compound observations
  int<lower = 1, upper = J_comp> comp[N_comp]; // compound index
  int<lower = 0> K_comp;  // number of compound-level predictors
  matrix[N_comp, K_comp] X_comp;  // compound-level predictors

// hw sample data
  int<lower = 0> N_hw;  // number of compound observations
  int<lower = 0, upper = 1> y_hw_hf[N_hw];  // hw HF183 observations
  // int<lower = 0, upper = 1> y_hw_mn[N_hw];  // no hw Mnif observations
  int<lower = 1, upper = J_comp> comp_hw[N_hw]; // hw compound index
  int<lower = 0> K_hw;  // number of hw sample-level predictors
  matrix[N_hw, K_hw] X_hw;  // hw predictors

// ds sample data
  int<lower = 0> N_ds;  // number of compound observations
  int<lower = 0, upper = 1> y_ds_hf[N_ds];  // ds HF183 observations
  int<lower = 0, upper = 1> y_ds_mn[N_ds];  // ds Mnif observations
  int<lower = 1, upper = J_comp> comp_ds[N_ds]; // ds compound index
  int<lower = 0> K_ds;  // number of ds sample-level predictors
  matrix[N_ds, K_ds] X_ds;  // ds predictors

// ls sample data
  int<lower = 0> N_ls;  // number of compound observations
  int<lower = 0, upper = 1> y_ls_hf[N_ls];  // ls HF183 observations
  int<lower = 0, upper = 1> y_ls_mn[N_ls];  // ls Mnif observations
  int<lower = 1, upper = J_comp> comp_ls[N_ls]; // ls compound index
  int<lower = 0> K_ls;  // number of ls sample-level predictors
  matrix[N_ls, K_ls] X_ls;  // ls predictors

// sens/spec data
  int<lower = 0> J_spec_hf;
  int<lower = 0> y_spec_hf[J_spec_hf];
  int<lower = 0> n_spec_hf[J_spec_hf];
  int<lower = 0> J_sens_hf;
  int<lower = 0> y_sens_hf[J_sens_hf];
  int<lower = 0> n_sens_hf[J_sens_hf];
  int<lower = 0> J_spec_mn;
  int<lower = 0> y_spec_mn[J_spec_mn];
  int<lower = 0> n_spec_mn[J_spec_mn];
  int<lower = 0> J_sens_mn;
  int<lower = 0> y_sens_mn[J_sens_mn];
  int<lower = 0> n_sens_mn[J_sens_mn];

// prior predictive check?
```

```
  int<lower = 0, upper = 1> prior_only;
}

parameters{
// compound prevalence linear model parameters
  vector[K_comp] b_comp; // compound-level coefficients
  real mu_comp;  // mean of compound-varying intercept
  real<lower = 0> sigma_comp;  // SD of compound-varying intercept
  vector<offset = mu_comp, multiplier = sigma_comp>[J_comp] a_comp;

// sample-level parameters
  vector[K_hw] g_hw;  // hw predictor coefficients
  vector[K_ds] g_ds;  // ds predictor coefficients
  vector[K_ls] g_ls;  // ls predictor coefficients
  real<lower = 0> sigma_type;  // SD of sample differences
  vector<multiplier = sigma_type>[N_type] a_type;  // sample type-varying intercept

// sens/spec meta-analysis parameters
  real mu_logit_spec_hf;
  real mu_logit_sens_hf;
  real mu_logit_spec_mn;
  real mu_logit_sens_mn;
  real<lower = 0> sigma_logit_spec_hf;
  real<lower = 0> sigma_logit_sens_hf;
  real<lower = 0> sigma_logit_spec_mn;
  real<lower = 0> sigma_logit_sens_mn;
  vector<offset = mu_logit_spec_hf, multiplier = sigma_logit_spec_hf>[J_spec_hf]
logit_spec_hf;
  vector<offset = mu_logit_sens_hf, multiplier = sigma_logit_sens_hf>[J_sens_hf]
logit_sens_hf;
  vector<offset = mu_logit_spec_mn, multiplier = sigma_logit_spec_mn>[J_spec_mn]
logit_spec_mn;
  vector<offset = mu_logit_sens_mn, multiplier = sigma_logit_sens_mn>[J_sens_mn]
logit_sens_mn;
}

transformed parameters{
  vector[J_spec_hf] spec_hf = inv_logit(logit_spec_hf);
  vector[J_sens_hf] sens_hf = inv_logit(logit_sens_hf);
  vector[J_spec_mn] spec_mn = inv_logit(logit_spec_mn);
  vector[J_sens_mn] sens_mn = inv_logit(logit_sens_mn);
}

model{
// linear model for compound contamination
  vector[N_comp] logit_p_comp = a_comp[comp] + X_comp * b_comp;

// linear models for sample-type specific prevalence
  vector[N_hw] p_hw = inv_logit(logit_p_comp[comp_hw] + a_type[1] + X_hw * g_hw);
  vector[N_ds] p_ds = inv_logit(logit_p_comp[comp_ds] + a_type[2] + X_ds * g_ds);
  vector[N_ls] p_ls = inv_logit(logit_p_comp[comp_ls] + a_type[3] + X_ls * g_ls);

// adjust for sens/spec
  vector[N_hw] p_hw_hf = sens_hf[1] * p_hw + (1 - spec_hf[1]) * (1 - p_hw);
  vector[N_hw] p_hw_mn = sens_mn[1] * p_hw + (1 - spec_mn[1]) * (1 - p_hw);
  vector[N_ds] p_ds_hf = sens_hf[1] * p_ds + (1 - spec_hf[1]) * (1 - p_ds);
  vector[N_ds] p_ds_mn = sens_mn[1] * p_ds + (1 - spec_mn[1]) * (1 - p_ds);
```

```
  vector[N_ls] p_ls_hf = sens_hf[1] * p_ls + (1 - spec_hf[1]) * (1 - p_ls);
  vector[N_ls] p_ls_mn = sens_mn[1] * p_ls + (1 - spec_mn[1]) * (1 - p_ls);

// Likelihoods
  if(prior_only == 0){
  // samples
    y_hw_hf ~ binomial(1, p_hw_hf);
    // y_hw_mn ~ binomial(1, p_hw_mn); no Mnif for HW samples
    y_ds_hf ~ binomial(1, p_ds_hf);
    y_ds_mn ~ binomial(1, p_ds_mn);
    y_ls_hf ~ binomial(1, p_ls_hf);
    y_ls_mn ~ binomial(1, p_ls_mn);

  // validation studies
    y_spec_hf ~ binomial(n_spec_hf, spec_hf);
    y_sens_hf ~ binomial(n_sens_hf, sens_hf);
    y_spec_mn ~ binomial(n_spec_mn, spec_mn);
    y_sens_mn ~ binomial(n_sens_mn, sens_mn);
  }

// sample priors
  // compound-level
  a_comp ~ normal(mu_comp, sigma_comp);
  mu_comp ~ normal(0, 1.5);
  sigma_comp ~ normal(0, 0.5);
  b_comp ~ normal(0, 0.5);

  // sample-level
  a_type ~ normal(0, sigma_type);
  sigma_type ~ normal(0, 0.5);
  g_hw ~ normal(0, 0.5);
  g_ds ~ normal(0, 0.5);
  g_ls ~ normal(0, 0.5);

// validation priors
  logit_spec_hf ~ normal(mu_logit_spec_hf, sigma_logit_spec_hf);
  logit_sens_hf ~ normal(mu_logit_sens_hf, sigma_logit_sens_hf);
  sigma_logit_spec_hf ~ normal(0, .5);
  sigma_logit_sens_hf ~ normal(0, .5);
  mu_logit_spec_hf ~ normal(.5, 1);
  mu_logit_sens_hf ~ normal(.5, 1);
  logit_spec_mn ~ normal(mu_logit_spec_mn, sigma_logit_spec_mn);
  logit_sens_mn ~ normal(mu_logit_sens_mn, sigma_logit_sens_mn);
  sigma_logit_spec_mn ~ normal(0, .5);
  sigma_logit_sens_mn ~ normal(0, .5);
  mu_logit_spec_mn ~ normal(.5, 1);
  mu_logit_sens_mn ~ normal(.5, 1);
}

generated quantities{
// simulated sample outcome containers
  int y_hw_hf_sim[N_hw];
  int y_hw_mn_sim[N_hw];
  int y_ds_hf_sim[N_ds];
  int y_ds_mn_sim[N_ds];
  int y_ls_hf_sim[N_ls];
  int y_ls_mn_sim[N_ls];
```

```
  int n_pos_hw_hf;
  int n_pos_hw_mn;
  int n_pos_ds_hf;
  int n_pos_ds_mn;
  int n_pos_ls_hf;
  int n_pos_ls_mn;
  real<lower = 0, upper = 1> p_avg_comp;
  real<lower = 0, upper = 1> p_avg_hw;
  real<lower = 0, upper = 1> p_avg_ds;
  real<lower = 0, upper = 1> p_avg_ls;
  real<lower = 0, upper = 1> p_samp_avg_hw_hf;
  real<lower = 0, upper = 1> p_samp_avg_hw_mn;
  real<lower = 0, upper = 1> p_samp_avg_ds_hf;
  real<lower = 0, upper = 1> p_samp_avg_ds_mn;
  real<lower = 0, upper = 1> p_samp_avg_ls_hf;
  real<lower = 0, upper = 1> p_samp_avg_ls_mn;

// simulate type-specific probabilities
  vector[N_comp] logit_p_comp_sim = a_comp[comp] + X_comp * b_comp;
  vector[N_comp] p_comp_sim = inv_logit(logit_p_comp_sim);
  vector[N_hw] p_hw_sim = inv_logit(logit_p_comp_sim[comp_hw] + a_type[1] + X_hw *
g_hw);
  vector[N_ds] p_ds_sim = inv_logit(logit_p_comp_sim[comp_ds] + a_type[2] + X_ds *
g_ds);
  vector[N_ls] p_ls_sim = inv_logit(logit_p_comp_sim[comp_ls] + a_type[3] + X_ls *
g_ls);
// adjust simulations for sens/spec
  vector[N_hw] p_hw_hf_sim = sens_hf[1] * p_hw_sim + (1 - spec_hf[1]) * (1 -
p_hw_sim);
  vector[N_hw] p_hw_mn_sim = sens_mn[1] * p_hw_sim + (1 - spec_mn[1]) * (1 -
p_hw_sim);
  vector[N_ds] p_ds_hf_sim = sens_hf[1] * p_ds_sim + (1 - spec_hf[1]) * (1 -
p_ds_sim);
  vector[N_ds] p_ds_mn_sim = sens_mn[1] * p_ds_sim + (1 - spec_mn[1]) * (1 -
p_ds_sim);
  vector[N_ls] p_ls_hf_sim = sens_hf[1] * p_ls_sim + (1 - spec_hf[1]) * (1 -
p_ls_sim);
  vector[N_ls] p_ls_mn_sim = sens_mn[1] * p_ls_sim + (1 - spec_mn[1]) * (1 -
p_ls_sim);

// simulate sample observations
  y_hw_hf_sim = binomial_rng(1, p_hw_hf_sim);
  y_hw_mn_sim = binomial_rng(1, p_hw_mn_sim);
  y_ds_hf_sim = binomial_rng(1, p_ds_hf_sim);
  y_ds_mn_sim = binomial_rng(1, p_ds_mn_sim);
  y_ls_hf_sim = binomial_rng(1, p_ls_hf_sim);
  y_ls_mn_sim = binomial_rng(1, p_ls_mn_sim);

// summarize simulated samples
  n_pos_hw_hf = sum(y_hw_hf_sim);
  n_pos_hw_mn = sum(y_hw_mn_sim);
  n_pos_ds_hf = sum(y_ds_hf_sim);
  n_pos_ds_mn = sum(y_ds_mn_sim);
  n_pos_ls_hf = sum(y_ls_hf_sim);
  n_pos_ls_mn = sum(y_ls_mn_sim);

// mean prevalence predictions
```

```
  p_avg_comp = mean(p_comp_sim);
  p_avg_hw = mean(p_hw_sim);
  p_avg_ds = mean(p_ds_sim);
  p_avg_ls = mean(p_ls_sim);
  p_samp_avg_hw_hf = mean(p_hw_hf_sim);
  p_samp_avg_hw_mn = mean(p_hw_mn_sim);
  p_samp_avg_ds_hf = mean(p_ds_hf_sim);
  p_samp_avg_ds_mn = mean(p_ds_mn_sim);
  p_samp_avg_ls_hf = mean(p_ls_hf_sim);
  p_samp_avg_ls_mn = mean(p_ls_mn_sim);
}
```

(5) Johnston, C.; Ufnar, J. a; Griffith, J. F.; Gooch, J. a; Stewart, J. R. A Real-Time QPCR Assay for the Detection of the NifH Gene of Methanobrevibacter Smithii, a Potential

Indicator of Sewage Pollution. *Journal of applied microbiology* **2010**, *109* (6), 1946–1956. https://doi.org/10.1111/j.1365-2672.2010.04824.x.

(25) McElreath, R. *Statistical Rethinking: A Bayesian Course with Examples in R and Stan*, 2nd ed.; CRC texts in statistical science; Taylor and Francis, CRC Press: Boca Raton, 2020.
